# A single cell framework for multi-omic analysis of disease identifies malignant regulatory signatures in mixed phenotype acute leukemia

**DOI:** 10.1101/696328

**Authors:** Jeffrey M. Granja, Sandy Klemm, Lisa M. McGinnis, Arwa S. Kathiria, Anja Mezger, Benjamin Parks, Eric Gars, Michaela Liedtke, Grace X.Y. Zheng, Howard Y. Chang, Ravindra Majeti, William J. Greenleaf

**Author notes:** These authors contributed equally to this study (Jeffrey M. Granja, Sandy Klemm, Lisa M. McGinnis). Correspondence to: W.J.G., S.K., L.M.M.

## Abstract

In order to identify the molecular determinants of human diseases, such as cancer, that arise from a diverse range of tissue, it is necessary to accurately distinguish normal and pathogenic cellular programs.^1–3^ Here we present a novel approach for single-cell multi-omic deconvolution of healthy and pathological molecular signatures within phenotypically heterogeneous malignant cells. By first creating immunophenotypic, transcriptomic and epigenetic single-cell maps of hematopoietic development from healthy peripheral blood and bone marrow mononuclear cells, we identify cancer-specific transcriptional and chromatin signatures from single cells in a cohort of mixed phenotype acute leukemia (MPAL) clinical samples. MPALs are a high-risk subtype of acute leukemia characterized by a heterogeneous malignant cell population expressing both myeloid and lymphoid lineage-specific markers.^4, 5^ Our results reveal widespread heterogeneity in the pathogenetic gene regulatory and expression programs across patients, yet relatively consistent changes within patients even across malignant cells occupying diverse portions of the hematopoietic lineage. An integrative analysis of transcriptomic and epigenetic maps identifies 91,601 putative gene-regulatory interactions and classifies a number of transcription factors that regulate leukemia specific genes, including *RUNX1*-linked regulatory elements proximal to *CD69*. This work provides a template for integrative, multi-omic analysis for the interpretation of pathogenic molecular signatures in the context of developmental origin.

## Main

To identify pathologic features within neoplastic cells, we first aimed to establish molecular features of normal development for comparison. Since MPALs present with features of multiple hematopoietic lineages, we first constructed independent immunophenotypic, transcriptomic and epigenetic maps of normal blood development using droplet-based CITE-seq^6^ (single-cell antibody derived tag and RNA sequencing) and single-cell ATAC-seq (scATAC-seq, single-cell chromatin accessibility profiling)^7^ on bone marrow and peripheral blood mononuclear cells (**Figure 1a**). For CITE-seq analyses, we simultaneously generated 10x Genomics 3^’^ single-cell RNA sequencing^8^ (scRNA-seq) and antibody derived tag sequencing^6^ (scADT-seq) libraries from 35,882 bone marrow mononuclear cells (BMMCs, n = 12,602), CD34^+^ enriched BMMCs (n = 8,176), and peripheral blood mononuclear cells (PBMC, n = 14,804). On average, 1,273 informative genes (2,370 unique transcript molecules) were detected per cell and replicates were highly correlated (Supplementary Figure 1a-b). We then selected a feature set of transcripts to mitigate batch effects and linearly projected retained transcript counts into a lower dimensional space using Latent Semantic Indexing (LSI, see Online Methods).^9, 10^ Cells were clustered using Seurat’s Shared Nearest Neighbor approach^11^, annotated using a manually curated maker gene list, and visualized using uniform manifold approximation and projection (UMAP)^12^ (**Figure 1b**, Supplementary Figure 1c-d).

**Figure 1.**
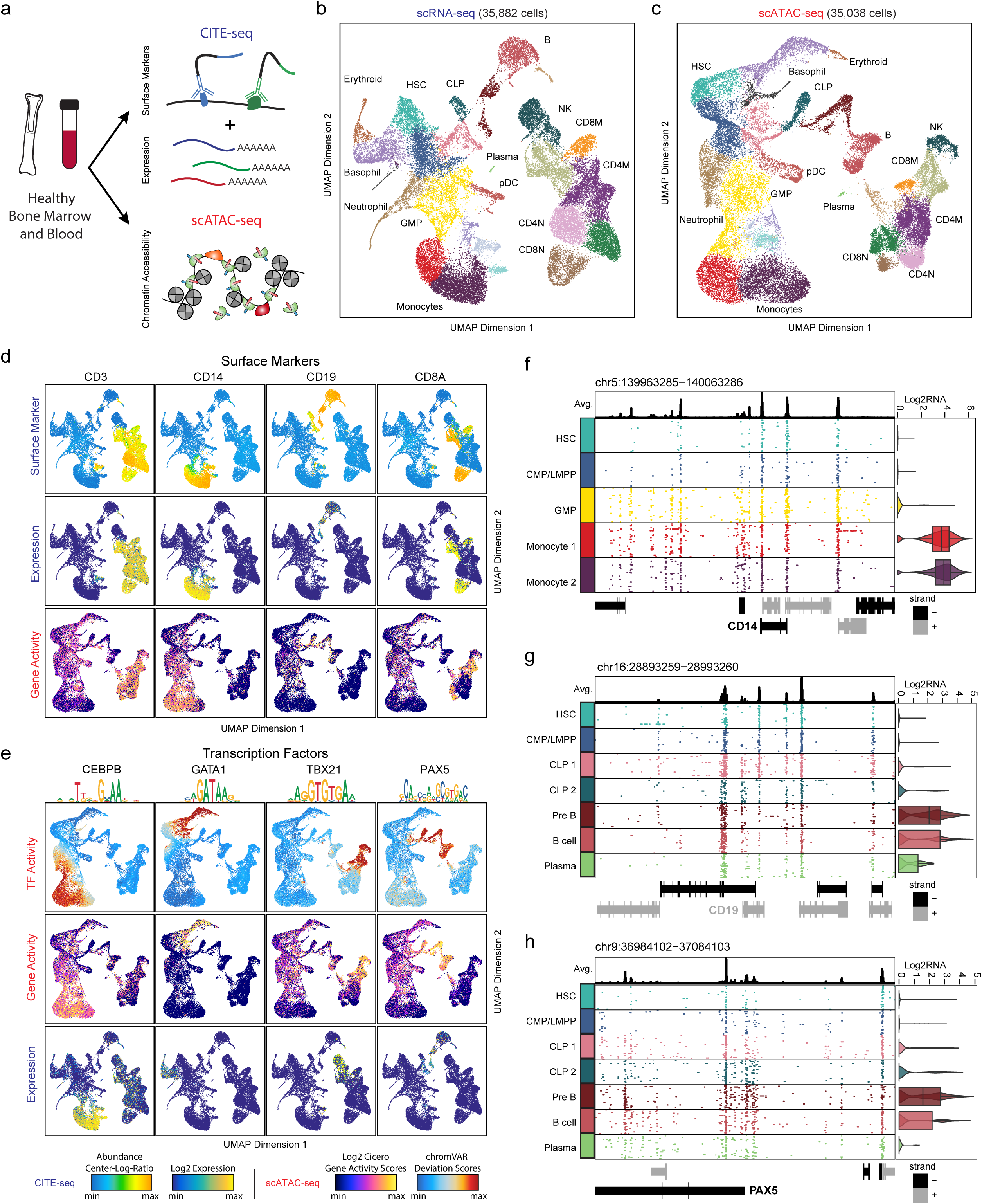
Multi-omic epigenetic and phenotypic analysis of human hematopoiesis. **a**, Schematic of multi-omic profiling of chromatin accessibility, transcription, and cell surface antibody abundance on healthy bone marrow and peripheral blood mononuclear cells using scATAC-seq and CITE-seq (combined single-cell RNA and antibody derived tag sequencing). **b**, scRNA-seq LSI UMAP projection of 35,882 single cells of healthy hematopoiesis. **c**, scATAC-seq LSI UMAP projection of 35,038 single cells of healthy hematopoiesis. **d**, Surface marker overlay on single-cell RNA UMAP (**b**) of (Top) ADT antibody signal (CLR normalized), (Middle) single-cell RNA, and (Bottom) log2 gene activity scores for *CD3*, *CD14*, *CD19*, and *CD8A*. **e**, Transcription factor overlay on single-cell ATAC UMAP (**c**) of (Top) TF deviations, (Middle) gene activity scores, and (Bottom) single-cell RNA for *CEBPB*, *GATA1*, *TBX21*, and *PAX5*. **f-h**, Multi-omic tracks; (Top) average track of all clusters displayed, (Middle) binarized 100 random scATAC-seq tracks for each locus at 100bp resolution and (right) scRNA-seq log2 distribution of normalized expression for each cluster. **f,** Multi-omic track of *CD14* (specific in these clusters for monocytes) across monocyte development from HSC progenitor cells. **g,** Multi-omic track of *CD19* (specific in these clusters for pre B cells) across B cell development. **h,** Multi-omic track of *PAX5* (specific in these clusters for pre B cells) across B cell development.

We next established an epigenetic map of normal hematopoiesis by measuring chromatin accessibility across 35,038 single BMMCs (n = 16,510), CD34^+^ BMMCs (n = 10,160), and PBMCs (n = 8,368) using droplet scATAC-seq (10x Genomics)^7^. These cells exhibited a canonical fragment size distribution with clearly resolved sub-, mono-, and multi-nucleosomal modes, a high signal-to-noise ratio at transcription start sites, an average of 11,597 uniquely accessible fragments per cell on average, and a majority (61%) of Tn5 insertions aligning within peaks (Supplementary Figure 2a-c). After pooling all scATAC-seq profiles from each experiment, we confirmed higher reproducibility across replicates than across different samples, similar to the scRNA-seq analysis (Supplementary Figure 2d). Using LSI, Seurat’s Shared Nearest Neighbor clustering, and UMAP, we generated a chromatin accessibility map of hematopoiesis that complements the transcriptional map of hematopoiesis (**Figure 1c**, Supplementary 2e-f).

To validate the proposed transcriptomic and epigenetic single-cell maps of hematopoiesis, we directly visualized lineage-restricted cell-surface marker and transcription factor enrichment across each map. As anticipated, both scADT- and scRNA-seq measurements of surface makers demonstrate *CD3* enrichment across bone marrow and peripheral T cells; *CD14* enrichment within the monocytic lineage; broad up regulation of *CD19* across the B cell lineage; and *CD8A* enrichment within cytotoxic T lymphocytes (**Figure 1d**)^13^. Estimates of gene activity based on correlated variation in promoter and distal peak accessibility (Cicero^14^) broadly recapitulates this pattern, confirming that lineage specification is consistently reflected across the phenotypic, transcriptional and epigenetic maps of hematopoietic development (**Figure 1d**). We then visualized our high quality scADT-seq using UMAP and found that we could broadly recapitulate our transcriptomic hematopoietic map (Supplementary Figure 3a-d). To further support these cell type identifications and developmental mappings, we show concordance between three separate single-cell measurements, including direct transcript measurements from the scRNA-seq dataset, inferred gene activity scores from the scATAC-seq dataset, and TF activity using chromVAR^15^, for key developmental transcription factors, including *CEBPB* in monocytic development, *GATA1* within the erythroid lineage, and *TBX21* in NK and CD8^+^ T memory cells, and *PAX5* in B cell and plasmacytoid dendritic cell development (**Figure 1e**). High-resolution single cell multi-omic tracks for key marker genes in each of the identified lineages further support these identifications (**Figure 1f-h,** Supplementary Figure 4a-h). Collectively these results show that the proposed multi-omic maps of healthy hematopoiesis are consistent and broadly capture essential phenotypic, transcriptomic and epigenetic features of blood development.

Recent work has shown that immunophenotypically-distinct subpopulations of MPAL blasts share similar genomic lesions within a patient, and that cells from one lineage can reconstitute the alternate lineage in xenograft models^16^, suggesting that MPAL lineage plasticity may be epigenetically regulated. To explore the nature of this regulatory and phenotypic dysfunction, we assayed six MPAL samples including three T-myeloid (T/M) MPALs (MPAL1-3), 1 B-myeloid (B/M) MPAL (MPAL4), and one T/M MPAL sampled before CALGB chemotherapy (MPAL5) and after post-treatment relapse (MPAL5R) (see Supplementary Table 1). Across these samples, we observed extensive immunophenotypic heterogeneity (via diagnostic flow cytometric analysis) including bilineal patterns (multiple blast populations expressing both lymphoid and myeloid lineage antigens), biphenotypic patterns (a dominant blast population that simultaneously expresses both lymphoid and myeloid antigens), and both patterns (Supplementary Figures 5a-c, 6a-c). We then performed Whole Exome Sequencing (WES) and found mutational profiles similar to previous studies (Supplementary Figure 6d)^16, 17^. To further profile our MPAL samples, we performed CITE-seq (18,056 cells) and scATAC-seq (35,423 cells) on either peripheral blood or bone marrow aspirates from these MPAL patients, observing similar high data quality to that obtained for healthy samples (Supplementary Figure 7a-f).

Using our transcriptomic and chromatin landscapes of healthy hematopoiesis, we next sought to develop an analytical framework to identify the hematopoietic developmental signature at single-cell resolution. First, the chromatin and gene expression signatures of single cells are projected into our ATAC- and RNA-based healthy hematopoietic map’s LSI subspace, and the results are then visualized using UMAP (**Figure 2a**, Supplementary Figure 8a). Next, by determining the closest hematopoietic cells to the projected cells we can identify the hematopoietic developmental compartment. This method does not require defining discrete cell type boundaries and uses a large feature set to robustly position cells within the continuous landscape of hematopoiesis. To validate this approach, we first projected downsampled, published bulk RNA-seq and ATAC-seq data^18^ from FACS-sorted subpopulations into our chromatin and transcription hematopoietic maps and found high concordance with our healthy hematopoietic map and cluster definitions (Supplementary Figure 8b). To further validate our approach, we projected published scRNA-seq^19^ and scATAC-seq^20–22^ data from different platforms and different genomes on our chromatin and transcription hematopoietic maps and found striking agreement (Supplementary Figure 8c). These results confirm that this method can accurately identify the hematopoietic signature for chromatin and gene expression at single-cell resolution.

**Figure 2.**
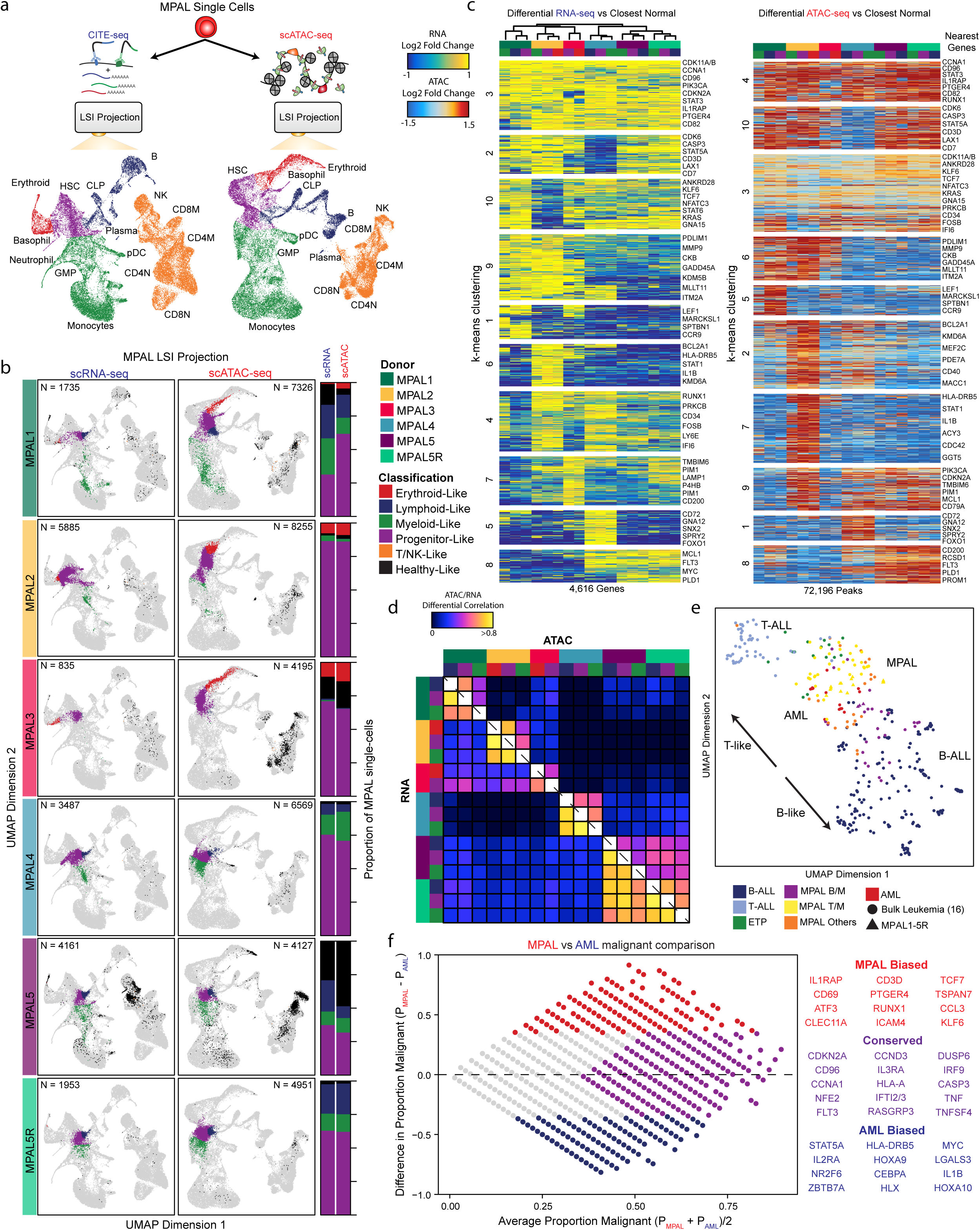
Multi-omic projection of MPALs into hematopoiesis identifies normal and leukemic programs. **a**, Schematic for projection of MPAL single cells onto hematopoiesis for both scRNA-seq and scATAC-seq classified into broad hematopoietic compartments. **b,** (Left) MPAL single cell projections into hematopoiesis for both scRNA-seq and scATAC-seq. (Right) The proportion of MPAL cells that were broadly classified as healthy or disease and their respective hematopoietic compartment. **c,** (Left) scRNA-seq heatmap of up-regulated genes log2 fold changes comparing MPAL disease subpopulations to closest non-redundant normal cells. Differential genes were clustered with k-means (k=10) based on their log2 fold changes. (Right) scATAC-seq heatmap of differentially up-regulated accessible peaks log2 fold changes comparing MPAL disease subpopulations to closest non-redundant normal cells. Differential peaks were clustered with k-means (k=10) based on their log2 fold changes. **d,** Pearson correlation of differentially up-regulated genes and peaks across all MPAL subpopulations. **e,** LSI UMAP of differentially up-regulated gene expression profiles across bulk leukemias^16^ and MPAL samples assayed in this study, colored by WHO 2016 classifications^5^. **f,** (Left) MA plot comparing the proportion of malignant (up-regulated) gene expression profiles in AML and MPALs. The x-axis represents for each up-regulated gene, the average proportion of AML and MPAL patient subpopulations broadly up-regulated (LFC > 0.5). The y-axis represents for each up-regulated gene, the difference in the proportion of MPAL and AML patient subpopulations up-regulated (LFC > 0.5). (Right) Genes that are more malignant biased to AMLs, MPALs and conserved across both AMLs and MPALs.

Using this LSI projection framework and landscapes of healthy hematopoiesis, we next sought to deconvolve the normal and leukemic signatures of MPAL samples at single-cell resolution. First, the leukemic single cells are projected into the hematopoietic linear LSI subspace. Next we identify a non-redundant set of healthy hematopoietic cells that were nearest neighbor normal cells to each leukemic cell, irrespective of their cell-type boundaries. Lastly, we compute the differences between the leukemic cells and nearest normal cells to identify the leukemic specific signature. We first tested our approach by analyzing recently published scRNA-seq data from acute myeloid leukemia (AML) patient samples^19^. By projecting the AMLs into our healthy hematopoietic map, we see general agreement with previous classifications without need for the establishment of potential arbitrary cell-type boundaries on normal hematopoiesis (Supplementary Figure 9a-c). We next projected our phenotypically diverse MPAL patient samples onto our hematopoietic maps and discovered broad epigenetic and gene expression diversity. To further resolve this diversity, we grouped MPAL cells within individual patients into broad hematopoietic developmental compartments: progenitors-like (purple) comprising human stem cell and multipotent progenitor-like cells, lymphoid-like (blue) containing lymphoid-primed multipotent progenitors, erythroid-like (red) which include megakaryocyte-erythroid progenitors, myeloid-like (green) which include granulocyte-monocyte progenitors, and T/NK-like (orange) which include differentiated T and NK cells^23^ (**Figure 2a-b**). Strikingly, we see that the scADT-seq data clearly resolves the dominant MPAL subpopulations in MPAL1 and MPAL5; however it does not fully capture the transcriptional diversity of MPALs 2-4 (Supplementary Figure 10a). We visualized these projected MPALs colored by these broad hematopoietic compartments, observing the expected high concordance between the scRNA- and scATAC-seq classifications (**Figure 2b**). Comparing MPAL gene expression to this healthy nearest neighbor set allowed the identification of pathogenic differential gene expression for MPALs from different compartments. In total, we identified 4,616 genes that were significantly up-regulated (LFC > 0.5 and FDR < 0.01) in at least one MPAL subpopulation across the six patient samples, and grouped these genes with k-means clustering (**Figure 2c**). We further categorized the most conserved differential genes, TFs and KEGG pathways across the MPALs (Supplementary Figure 11a-c). Using the same approach for the scATAC-seq data, we performed differential peak testing for each MPAL subpopulation and found 72,196 significantly up-regulated peaks (LFC > 0.5 and FDR < 0.05) in at least one MPAL subpopulation (**Figure 2c**). Multi-omic differential tracks for the cyclin dependent kinase *CDK11A* and cyclin dependent kinase inhibitor *CDKN2A*, genes that are recurrently mutated in MPAL^16, 24^, demonstrate these leukemia-specific ATAC- and RNA-seq differences (Supplementary Figure 11d-e). Additionally, we calculated Pearson correlations of the differential genes and peaks; and found that transcription and accessibility differs significantly *across* patients, but is relatively conserved across subpopulations *within* patients. (**Figure 2d**).

To compare the MPAL hematopoietic compartments’ leukemic programs to previous studies, we downsampled bulk leukemia RNA-seq and projected onto our transcriptomic hematopoietic UMAP for childhood AMLs, B-acute lymphoblastic leukemias (B-ALLs), early T-cell precursor T-acute lymphoblastic leukemias (ETP T-ALLs), non-ETP T-ALLs and MPALs^16^ (Supplementary Figure 12a-b). We then calculated differential expression with respect to the closest normal cell populations to identify their respective leukemic programs. Next, we performed LSI on variable malignant genes across all the leukemia subtypes, including MPALs 1-5, and then visualized these patients with UMAP (**Figure 2e,** Supplementary Figure 12c-d). Interestingly, we found large differences in the leukemic programs across various leukemias including T-ALLs, B-ALLs, and across different cytogenetic subtypes. In addition, we found that the MPALs assayed in this study were representative of previous characterized MPALs^16^ (**Figure 2e**). Given that we were insufficiently powered to detect unique leukemic differences between AML and our MPAL samples when analyzing downsampled bulk data, we compared the malignant transcriptomic profiles identified from re-analyzing AML scRNA-seq data^18^ with our MPALs in order to dissect further these unique malignant signatures (**Figure 2c**, Supplementary Figure 9c). To this end, we identified genes that were more commonly universally upregulated in AMLs, in MPALs, or jointly upregulated in both leukemias (**Figure 2f**, Supplementary Figure 9c). These gene sets provide fine-grained phenotypic resolution comparing the differences and similarities between AML and MPAL leukemic programs and suggest possible insight into why MPALs respond poorly to AML treatment^25, 26^.

Having compared our leukemic transcriptomic programs to other studies we wanted to identify the key TFs that regulate these programs. First, we identified which TF were differentially enriched in each k-means cluster of differentially accessible peaks observed in Figure 2c. (**Figure 3a**). We found that *RUNX1* motifs were highly enriched in both cluster 4 and 10 – the two clusters corresponding to the most commonly shared accessible elements across MPAL subset populations. In addition, *RUNX1* is significantly up-regulated in about half (7/17) of the MPAL subpopulations. *RUNX1* is one of the most frequently mutated genes across hematologic malignancies acting as both a tumor suppressor with loss-of-function mutations in AML^27^, myelodysplastic syndrome (MDS)^28^, and ETP T-ALL^29, 30^, and as a putative oncogene in non-ETP T-ALL^31, 32^. Furthermore, wildtype *RUNX1* has been implicated as a potential driver of leukemogenesis in core-binding factor (CBF) leukemia^33^ and mixed lineage leukemia^34^.

**Figure 3.**
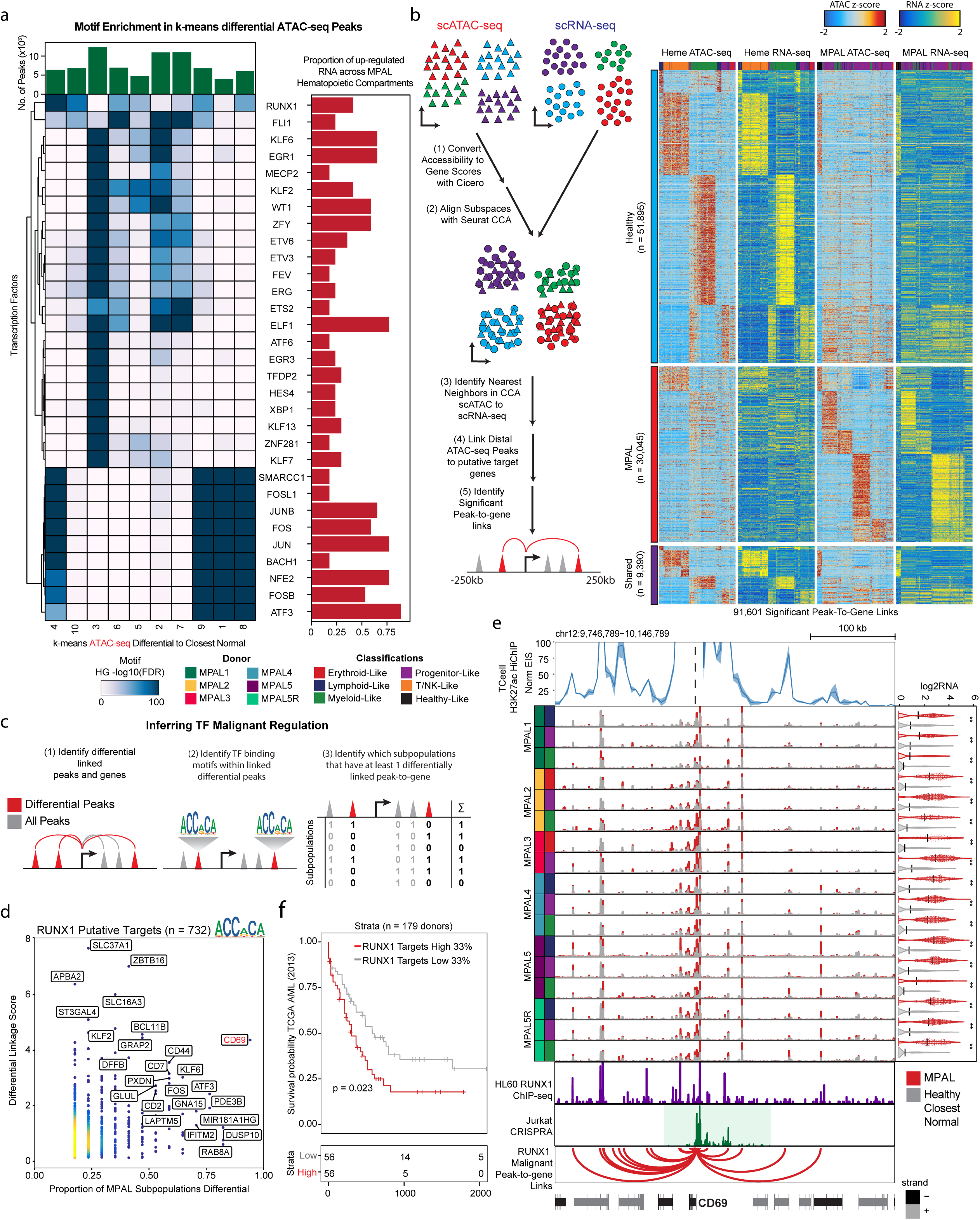
Integrative scATAC and scRNA-seq analyses nominate putative transcription factors that regulate leukemic programs. **a**, (Left) Hypergeometric TF motif enrichment FDR in differentially accessible peaks across each k-means clusters identified in Figure 2c. TFs are also identified as being differentially expressed and enriched in at least 3 MPAL hematopoietic compartments. (Top) Number of accessible peaks in each k-means cluster. (Right) Proportion of differentially up-regulated TF gene expression profiles across MPAL hematopoietic compartments. **b**, (Left) Schematic for alignment of scATAC and scRNA-seq data to link putative regulatory regions to target genes. First, scATAC-seq data is converted from accessible peaks to inferred gene activity scores using Cicero. Second, these gene activity scores and scRNA-seq expression are aligned into a common subspace using Seurat’s Canonical Correlation Analyses. Third, each scATAC-seq cell is assigned its nearest scRNA-seq neighbor. Fourth, ATAC-seq peaks within 2.5-250kb to a gene promoter are correlated within the healthy hematopoietic and MPAL knn groupings. Lastly, significant peak-to-gene links are identified by correlating peaks to genes on different chromosomes. (Right) Heatmaps of 91,601 peak-to-gene links across hematopoiesis and MPALs. (Top) peak-to-gene links that are identified only within hematopoiesis, (Middle) peak-to-gene links that are unique to MPALs, and (Bottom) peak-to-gene links identified in both hematopoiesis and MPALs. **c**, Schematic for identifying genes that are putatively regulated by the transcription factor of interest. **d**, *RUNX1* putative target genes differentially up-regulated in at least 1 MPAL subpopulations. The x-axis represents the proportion of MPAL subpopulations that are differential in both scRNA-seq and a linked accessible peak. The y-axis represents the cumulative linkage score between differentially up-regulated peaks linked to differentially up-regulated genes. **e,** *CD69* multi-omic differential track (Top) T cell Th17 K27ac HiChIP virtual4C of the *CD69* locus, shading represents standard deviation between biological replicates (n = 2). (Middle) Aggregated scATAC tracks showing MPAL disease subpopulations (red) and aggregated nearest-neighbor healthy (grey). (Right) Distribution of log2 normalized expression of *CD69* for MPAL disease subpopulations (red) and closest normal cells (grey); black line represents the mean and asterisk denote significance (LFC > 0.5 and FDR < 0.01). (Bottom) HL60 AML line ChIP-seq data across *CD69* locus, Jurkat CRISPRa tiling screen across the *CD69* locus and *RUNX1* identified malignant peak-to-gene links. **f**, Kaplan-Meier curve for TCGA AML patients (n=179) stratified by *RUNX1* putative target genes top 33% vs bottom 33% (p-value = 0.023).

To link *RUNX1* and other putative regulatory TFs to their leukemic programs we first developed an analytical framework that utilizes both our transcriptomic and chromatin single-cell data to link putative regulator peaks to target genes. Using our matched scATAC and scRNA data for all MPALs and concordant hematopoietic maps, and aligned each single-cell into a common subspace using Canonical Correlation Analyses (CCA)^10, 11, 35^. For each scATAC cell, we identified the nearest scRNA-seq neighbor (**Figure 3b**, Supplementary Figure 13a-b). We found that the mapping of scATAC cell clusters to scRNA-defined cell clusters were highly consistent (single-cell overlap of 52% across 26 clusters) (Supplementary Figure 14a-d). We then aggregated our scATAC cells based on nearest neighbors in the LSI subspace using Cicero^14^ and created a corresponding scRNA aggregate for each cluster using the constructed CCA alignment. We next identified 91,601 peak-to-gene links by correlating accessibility changes of ATAC peaks within 250 kb of the gene promoter with the expression of the gene independently for both healthy and MPAL aggregates (**Figure 3b**). This analysis revealed peak-to-gene links that were specific to healthy hematopoiesis, others that were specific to MPALs, and a conserved subset that was shared across both hematopoiesis and MPALs. We hypothesize that the MPAL-specific peak-to-gene links may be important for leukemic gene regulation. Overall, the identified set of peak-to-gene links had similar distributions for peaks mapped per gene, genes mapped per peak, number of skipped genes and the peak-to-gene as previously observed in a similar linkage analyses^2^ (Supplementary Figure 14e). To further support these peak-to-gene links, we used previously published K27ac HiChIP in primary T cells and a Human Coronary Artery Smooth Muscle Cells (HCASMC) cell-line and found that the T/NK biased peak-to-gene links were more enriched in the T cells than the HCASMC cell line^36^ (Supplementary Figure 14f). We next examined GTEx eQTL mappings within our inferred peak-to-gene links, finding enrichment of eQTLs in several functionally related categories such as Whole Blood and Lymphocytes (Supplementary Figure 14g). To demonstrate the utility of these peak-to-gene links, we linked differentially accessible regions to known leukemic genes such as the surface protein *CD96,* the leukemic stem cell marker *IL1RAP*, the cytokine receptor *FLT3*, and apoptosis regulator *MCL1* (Supplementary Figure 15a-d). Overall, these analyses, support that our peak-to-gene links are highly enriched in immune regulation and across other previously published linkage data sets^2, 36^.

Having established a high-quality set of peak-to-gene links, we aimed to identify the set of malignant genes putatively regulated by *RUNX1*. First, we utilized our peak-to-gene links to identify differential peaks linked to a differential gene within at least 2 MPAL subpopulations. Next, we selected all linked differential accessibility sites that contain the *RUNX1* motif. Finally, for each linked gene we combined all linked peaks to create a differential linkage score (see methods) and compared this score to the proportion of MPAL subpopulations that exhibited differential expression and accessibility in at least one linked peak and target gene (a measure of how common this *RUNX1*-driven dysfunction is across MPAL subsets) (**Figure 3c**). Using this approach, we found 732 genes putatively regulated by a RUNX1-containing distal element in at least 2 MPAL subsets, and found that *CD69*, gene implicated in lymphocyte activation through initiation of JAK/STAT signaling^37^ and lymphocyte retention in lymphoid organs^38^, was both highly enriched in the calculated differential linkage score and was observed to be differentially up-regulated in almost every MPAL subpopulation (**Figure 3d**). To further support *RUNX1* predicted regulation of *CD69*^39^, we incorporated T cell K27ac HiChIP^36^, CRISPRa screens^40^, and *RUNX1* ChIP-seq^41^ onto our multi-omic differential track. These orthogonal data sets show *RUNX1* binding to linked distal regulatory regions (**Figure 3e**). Finally, by using the 732 identified *RUNX1* target genes to stratify TCGA AML^42^ patients by expression, we observe significantly decreased survival (p-value = 0.023) in donors with a high *RUNX1* target gene signature^42^ (**Figure 3f**). This analysis suggests that *RUNX1* is an important TF that putatively up-regulates a portion of the leukemic signature in MPAL and potentially AML.

Collectively, this work establishes an experimental and analytical approach for deconstructing cancer-specific features using integrative analysis of multiple single-cell technologies. We find that MPAL malignant programs are largely conserved across phenotypically heterogenous cells within individual patients; this observation is consistent with a previous report^16^ that MPAL cells likely originate from a multipotent progenitor cell, thereby sharing a common mutational landscape while populating different regions of the hematopoietic tree. We used integrative single-cell analyses to further define putative TF regulation of these malignant programs. We inferred that *RUNX1* acts as a potential oncogene in MPAL, regulating malignant genes associated with poor survival. We anticipate that similar approaches will be used in future studies to both identify the differentiation status of different tumor types (i.e. identify the closest “normal” cell type) as well as enable molecular dissection of molecular dysfunction in pathogenic cellular sub-types, with the ultimate goal of identifying personalized therapeutic targets through integrative single-cell molecular characterization.

## METHODS

### Experimental Methods

#### Description of Healthy Donors

PBMCs, BMMCs, and CD34+ bone marrow cells were obtained from healthy donors (AllCells).

#### Description of Leukemic Patients/Donors

Patient samples were collected with informed consent prospectively under a protocol approved by the Institutional Review Board (IRB) at Stanford University Medical Center (Stanford IRB, 42949, 18329, and 6453). Peripheral blood and bone marrow aspirate samples were processed by Lymphoprep (STEMCELL Technologies) gradient centrifugation and fresh frozen in Bambanker media. Diagnostic flow cytometric performed on bone marrow aspirate samples were analyzed. In all cases, a retrospective review of clinical parameters, hemogram data, peripheral blood smears, bone marrow aspirates, trephine biopsies, results of karyotype and flow cytometry studies was performed. Clinical follow-up information was obtained by retrospective review of the medical record charts. Cases were classified using the 2016 WHO classification of hematopoietic and lymphoid neoplasms^5^.

#### CITE-seq (combined single-cell antibody derived tag and RNA sequencing)

Combined single-cell RNA and antibody derived tag sequencing (CITE-seq) was performed as previously reported^6^ using the (version 2) Chromium Single Cell 3’ Library and Gel Bead Kit (Cat # 120237, 10X Genomics). Six thousand cells were targeted for each sample. Oligo-coupled antibodies were obtained from Biolegend indexed by PCR (10 cycles) with custom barcodes (see Supplementary Table 3), quantified by PCR using a PhiX Control v3 (Illumina, Cat #FC-110-3001) standard curve, and then sequenced on an Illumina NextSeq 550 together with scRNA-seq at no more than 60% of the total library composition (1.5pM loading concentration, 26 x 8 x 0 x 98 bp read configuration).

#### Single-cell ATAC-seq (scATAC-seq)

Single-cell ATAC-seq targeting four thousand cells per sample was performed using a beta-version of Chromium Single Cell ATAC Library and Gel Bead Kit (Cat # 1000110, 10X Genomics). Each sample library was uniquely barcoded and quantified by PCR using a PhiX Control v3 (Illumina, Cat #FC-110-3001) standard curve. Libraries were then pooled and loaded on a NextSeq 550 Illumina sequencer (1.4pM loading concentration, 33 x 8 x 16 x 33 bp read configuration) and sequenced to either 90% saturation or 30,000 unique reads per cell on average.

#### Whole-Exome Sequencing of Leukemic Patients/Donors

Genomic DNA was extracted from diagnostic peripheral blood mononuclear cells or bone marrow samples using Zymo Clean and Concentrator Kit. Library construction (Agilent SureSelect Human All Exon kit), quality assessment, and 150-bp paired-end sequencing (HiSeq4000) were performed by Novogene (Beijing, China). Reads with adapter contamination, uncertain nucleotides, and paired reads with >50% low-quality nucleotides were discarded. Paired-end reads were then aligned to the reference genome (GRCh37) using BWA software. Genome Analysis Toolkit (GATK) was used to ignore duplicates with Picard-tool. Filtered variants (SNP, INDELs) were identified using GATK HaplotypeCaller and variantFiltration. Variants obtained from initial analysis were further compared to dbSNP and 1000 Genomes database. Finally, missense, stopgain and frameshift mutations were compared against a custom panel of 300 genes that are recurrently mutated in hematologic malignancies as described previously^16, 17^.

### Analytical Methods

#### FACS Analysis

Flow cytometry was performed on a FACSCalibur or FACSCanto II (Becton Dickinson, San Jose, Ca, USA) cytometer using commercially available antibodies (Supplementary Table 2). Lymphocytes were identified by low side-scatter and bright CD45 expression. The gate was validated by backgating on CD3-positive or CD19-positive events. Blasts were identified by low side-scatter and dim CD45 expression. The gate was further assessed by backgating on CD34-positive events. Gates were drawn by additionally using isotype controls and internal positive and negative controls.

#### scADT-seq Analysis

Raw sequencing data were converted to fastq format using bcl2fastq (Illumina, version v2.20.0.422). ADTs were then assigned to individual cells and antibodies (see reference antibody barcodes in Supplementary Table 3) allowing for 2 and 3 barcode mismatches, respectively. Unique molecular counts for each cell and antibody were then generated by counting only barcodes with a unique molecular identifier. PBMC and BMMC ADT count data were transformed using the centered log ratio (CLR) as previously described^6^. PBMC and BMMC cells were visualized in two dimensions using uwot’s implementation of UMAP^43^ in R (n_neighbors = 50, min_dist = 0.4).

### scATAC-seq Analytical Methods

#### scATAC-seq Processing

Raw sequencing data was converted to fastq format using cellranger atac mkfastq (10x Genomics, version 1.0.0). Single-cell RNA-seq reads were aligned to the GRCh37 (hg19) reference genome and quantified using cellranger count (10x Genomics, version 1.0.0).

#### scATAC-seq Quality Control

To ensure that each single-cell was both adequately sequenced and had high signal to background, we filtered cells with less than 1000 unique fragments and enrichment at transcription start sites (TSS) was below 8. To calculate a TSS enrichment^2^, briefly Tn5 corrected insertions were aggregated +/- 2,000 bp relative (TSS strand-corrected) for each unique TSS genome wide. This profile was normalized to the mean accessibility +/- 1,900- 2,000 bp from the TSS, smoothed every 51bp, and the maximum smoothed value was reported as TSS enrichment in R. We estimate that the multiplet percentage for this study was around 4%^7^.

#### scATAC-seq Counts Matrix

To construct a counts matrix for each cell by each feature (window or peaks), we read each fragment.tsv.gz fill into a Genomic Ranges object. For each Tn5 insertion, the “start” and “end” of the ATAC-fragments, we used findOverlaps” to find all overlaps with the feature by insertions. Then we added a column with the unique id (integer) cell barcode to the overlaps object and fed this into a sparseMatrix in R. To calculate the fraction of reads/insertions in peaks, we used the colSums of the sparseMatrix and divided it by the number of insertions for each cell id barcode using “table” in R.

#### scATAC-seq Union Peak Set from Latent Semantic Indexing Clustering

We adapted a previous workflow for generating a union peak set that will account for diverse subpopulation structure^2, 9, 10^. First, we created 2.5kb windows genome wide using “tile(hg19chromSizes, width = 2500)” in R. Next, a cell by 2.5kb window sparse matrix was constructed as described above. The top 20,000 accessible windows were kept and the binarized matrix was transformed with the term frequency-inverse document frequency (“TF-IDF”) transformation^8^. Briefly we divided each index by the colSums of the matrix to compute the cell “term frequency”. Next we multiplied these values by log(1 + ncol(matrix) / rowSums(matrix)) which represents the “inverse document frequency”. This normalization resulted in a TF-IDF matrix that was then used as input to irlba’s singular value decomposition (SVD) implementation in R. The 2nd-25^th^ SVD dimensions (1^st^ dimension is correlated with cell read depth^15^) were used for creating a Seurat object and identified clusters using Seurat’s SNN graph clustering (v2.3.4) with “FindClusters” with a default resolution of 0.8. If the minimum cluster size was below 200 cells, the resolution was decreased until this criterion was reached leading to a final resolution of 0.8^N^ (where N represents the iterations until the minimum cluster size is 200 cells). For each cluster, peak calling was performed on Tn5-corrected insertions (each end of the Tn5-corrected fragments) using the MACS2 callpeak command with parameters “--shift −75 --extsize 150 --nomodel --call-summits --nolambda --keep-dup all -q 0.05.” The peak summits were then extended by 250bp on either side to a final width of 501bp, filtered by the ENCODE hg19 blacklist (https://www.encodeproject.org/annotations/ENCSR636HFF/), and then filtered to remove peaks that extend beyond the ends of chromosomes.

Overlapping peaks called were handled using an iterative removal procedure as previously described^2^. First, the most significant (MACS2 score) extended peak summit is kept and any peak that directly overlaps with that significant peak is removed. This process re-iterates to the next most significant peak until all peaks have either been kept or removed due to direct overlap with a more significant peak. The most significant 200,000 extend peak summits for each cluster were quantile normalized using “trunc(rank(v))/length(v)” in R (where v represents the vector of MACS2 peaks scores). These cluster peak sets were then merged and the previous iterative removal procedure was used. Lastly, we removed any peaks whose nucleotide content had any “N” nucleotides and any peaks mapping to chrY.

#### scATAC-seq-centric Latent Semantic Indexing clustering and visualization

scATAC-seq clustering was performed by adapting the strategy of Cusanovich et. al^9, 10^, to compute the term frequency-inverse document frequency (“TF-IDF”) transformation. Briefly we divided each index by the colSums of the matrix to compute the cell “term frequency.” Next, we multiplied these values by log(1 + ncol(matrix) / rowSums(matrix)), which represents the “inverse document frequency.” This resulted in a TF-IDF matrix that was used as input to irlba’s singular value decomposition (SVD) implementation in R. The first 50 SVD dimensions were used as input into a Seurat object and initial clustering was performed using Seurat’s (v2.3.4) SNN graph clustering “FindClusters” with a resolution of 1.5 (25 SVD dimensions for Healthy Hematopoiesis and 50 for Healthy Hematopoiesis and MPALs). We found that in some cases, that there was batch effect between experiments. To minimize this effect, we identified the top 50,000 variable peaks across the initial clusters (summed cell matrix for each cluster followed by edgeR logCPM transformation^44^). These 50,000 variable peaks were then used to subset the sparse binarized accessibility matrix and recomputed the “TF-IDF” transform. We used singular value decomposition on the TF-IDF matrix to generate a lower dimensional representation of the data by retaining the first 50 dimensions. We then used these reduced dimensions as input into a Seurat object and then final clusters were identified by using Seurat’s (v2.3.4) SNN graph clustering “FindClusters” with a resolution of 1.5 (50 SVD dimensions for Healthy Hematopoiesis and 50 for Healthy Hematopoiesis and MPALs). These same reduced dimensions were used as input to uwots implementation of UMAP (n_neighbors = 55, n_components = 2, min_dist = 0.45) and plotted in ggplot2 using R. We merged scATAC-seq clusters from a total of 36 clusters for hematopoiesis to 26 final clusters that best agreed with the scRNA-seq clusters (included in Supplemental Data). The objective of this analysis is to optimize feature selection, that minimizes batch effects, and enable projection of future data into the same manifold as described further below.

#### scATAC-seq Visualization in Genomic Regions

To visualize scATAC-seq data, we read the fragments into a GenomicRanges object in R. We then computed sliding windows across each region we wanted to visualize every 100 bp “slidingWindows(region,100,100)”. We computed a counts matrix for Tn5-corrected insertions as described above and then binarized this matrix. We then returned all non-zero indices (binarization) from the matrix (cell x 100bp intervals) and plotted them in ggplot2 in R with “geom_tile”. For visualizing aggregate scATAC-seq data, the binarized matrix above was summed and normalized. Scale factors were computed by taking the binarized sum in the global peakset and normalizing to 10,000,000. Tracks were then plotted in ggplot in R.

#### chromVAR

We measured global TF activity using chromVAR^15^. We used the cell by peaks and the CIS-BP motif (from chromVAR motifs “human_pwms_v1”) matches within these peaks from motifmatchr. We then computed the GC bias-corrected deviations using the chromVAR “deviations” function. We then computed the GC bias-corrected deviation scores using the chromVAR “deviationScores” function.

#### Gene Activity Scores using Cicero and Co-Accessibility

We calculated gene activities using the R package Cicero^14^. Briefly, we used the sparse binary cell by peaks matrix and created a cellDataSet, detectedGenes, and estimatedSizeFactors. We then created a “cicero_cds” with k=50 and the “reduced_coordinates” being the latent semantic indexing singular value decompositions coordinates (Hematopoiesis = 25, Hematopoiesis and MPALs = 50). This function returns aggregated accessibility across groupings of cells based on nearest-neighbor rules from FNN. We then identified all peak-peak linkages that were within 250 kb by resizing the peaks to 250 kb and 1bp and using “findOverlaps” in R. We calculated the pearson correlation for each unique peak-peak link and created a connections data.frame where the first column is peak_i and the second column is peak_j and third coaccessibility (pearson correlation). We then created a gene data.frame from the TxDb “TxDb.Hsapiens.UCSC.hg19.knownGene” in R. We then resized each gene from its TSS and created a window +/- 2.5 kb centered at the TSS and then annoted the “cicero_cds” using “annotate_cds_by_site”. We then calculated gene activities with “build_gene_activity_matrix” (coaccess cutoff of 0.35). Lastly we normalized the gene activities by using “normalize_gene_activities” and the read depth of the cells. We then log normalized these gene activities scores for interpretability by computing “log2(GA*1,000,000 +1)”.

### scRNA-seq Analytical Methods

#### scRNA-seq Processing

Raw sequencing data was converted to fastq format using cellranger mkfastq (10x Genomics, version 3.0.0). Single-cell RNA-seq reads were aligned to the GRCh37 (hg19) reference genome and quantified using cellranger count (10x Genomics, version 3.0.0). We kept genes that were present in both 10x gene transfer format (GTF) files v3.0.0 for hg19 and hg38 (https://support.10xgenomics.com/single-cell-gene-expression/software/release-notes/build). Mitochondrial and ribosomal genes were also filtered prior to further analysis. Genes remaining after these filtering steps we refer to as “informative” genes and enable cross genome comparison.

#### scRNA-seq Quality Control

We wanted to filter out cells whose transcripts were lowly captured and first plotted the distribution of genes detected and UMIs for all experiments. Based on these plots we chose to filter out cells that had less than 400 informative genes detected and 1000 UMIs. In addition, to lower multiplet representation, we filtered cells with above 10,000 UMIs. We estimate that the multiplet percentage for this study was around 6%^8^. We then plotted the correlation for each replicate experiment and found high reproducibility.

#### scRNA-seq-centric Latent Semantic Indexing clustering and visualization

We initially tested out a few methods for clustering scRNA but settled on an approach that enabled us to effectively capture the hematopoietic hierarchy without significant alteration of transcripts expression. We first log-normalized the transcript counts by first depth normalizing to 10,000 and adding a pseudo count prior to a log2 transform (log2(counts per ten thousand transcripts + 1)). Next, we identified the top 3000 variable genes and performed the TF-IDF transform on these 3000 genes. We then performed singular value decomposition (SVD) on this transformed matrix keeping the first 25 dimensions and used this as input to Seurat Shared Nearest Neighbor Clustering (v2.3.4) with an initial resolution of 0.2. We then summed the individual clusters single cells and computed the logCPM transformation, edgeR::cpm(mat,log=TRUE,prior.count=3), and then identified the top 2500 variable genes across these initial clusters. These variable genes were then used as input for a TF-IDF transform and then performed singular value decomposition (SVD) on this transformed matrix keeping the first 25 dimensions and used this as input to Seurat Shared Nearest Neighbor Clustering (v2.3.4) with an increased resolution of 0.6. We then summed the individual clusters single cells and computed the logCPM transformation, edgeR::cpm(mat,log=TRUE,prior.count=3), and then identified the top 2500 variable genes across these clusters. We then repeated this 1 more time (resolution 1.0) and then saved the final features and clusters. To align our clusters better with the scATAC-seq data we merged a total of 26 clusters from 31 initial clusters (included in Supplemental Data). These LSI dimensions were used as input to uwots implementation of UMAP (n_neighbors = 35, n_components = 2, min_dist = 0.45) and plotted in ggplot2 using R. The objective of this analysis is to optimize feature selection, that minimizes batch effects, and enable projection of future data into the same manifold as described further below.

### scATAC-seq and scRNA-seq Analytical Methods

#### LSI Projection for scATAC and scRNA-seq

We designed the above analytical approach to clustering single cell data because it optimized feature selection and enabled projection of new non-normalized data into low dimension manifold. To enable this analyses, when computing the TF-IDF transformation on the hematopoietic hierarchy, we kept the colSums, rowSums, and SVD from the previous run and then when projecting new data into this subspace, we first identified which row indices to zero out based on the initial TF-IDF rowSums. We then computed the “term frequency” by dividing by the colSums in these features. Next, we computed the “inverse document frequency” from the previous TF-IDF transform (diagonal(1+ncol(mat)/ rowSums(mat))) and computed the new TF-IDF transform. We then projected this TF-IDF matrix into the SVD subspace previous generated. To do this calculation, we computed the new coordinates by “t(TF_IDF) %*% SVD$u %*% diag(1/SVD$d)” where TF_IDF is the transformed matrix and SVD is the previous SVD run using irlba in R (3.5.1). We then computed the projected matrix by “SVD$u %*% diag(SVD$D) * t(V)” where V is the projected coordinates above. For projecting bulk RNA-seq, we downsampled previously published data to 5,000 reads in genes 100 times and then made a sparse matrix for projection as single cell data. For projecting bulk scATAC-seq, we downsampled previously published data to 10,000 reads in peaks 100 times and then made a binary sparse matrix for projection as single cell data.

#### Classification of MPAL single cells with scATAC and scRNA-seq

We wanted to classify MPAL single cells based on their disease state and hematopoietic progression. First, we determined which cells were healthy-like and disease-like. To do this analysis, we clustered all of the healthy hematopoietic cells with the MPAL of interest using our LSI workflow as described above (scRNA – 25 PCs, 1,000 variable genes and Seurat SNN resolution of 0.2, 0.8 and 0.8; scATAC - 25 PCs, 25,000 variable peaks and Seurat SNN resolution of 0.8 and 0.8). We then determined which clusters were “healthy-like” if a high percentage (>80% for scRNA, >90% for scATAC) of the cells were from the hematopoietic data. MPAL single cells belonging to these clusters were classified as “healthy-like” and the remaining disease-like. We note that we did not detect significant large-scale copy number amplifications with our previously described approach^7^, and the proportion of “disease-like” classified cells were consistent with our FACS estimation of percent blast cells. In order to accurately characterize these MPAL “disease-like” by their hematopoietic state, we established “hematopoietic compartments” across our scRNA and scATAC-seq maps that broadly characterized the hematopoietic continuum. The borders for these compartments were determined empirically using “fhs” in R, guided by the initial clusters and agreement across the scRNA and scATAC-seq classifications. After the hematopoietic continuum were classified, we then broadly classified the MPAL “disease-like” cells based on their projected nearest neighbor in the UMAP subspace. These classifications were used subsequently in differential analyses.

#### Identifying differential features with scATAC and scRNA-seq

To identify differential features for previously published AML data and MPALs, we constructed a nearest neighbor healthy aggregate using the following approach. First, we used FNN to identify the nearest 25 cells using “get.knnx(svdHealthy, svdProjected, k=25)” based on Euclidean distance between the projected cells and hematopoietic cells in LSI-SVD space. For each projected population, we used a minimum of 50 and maximum of 500 cells (random sampling) as input. Next, we took the unique of all hematopoietic single-cells and if this number was greater than 1.25 times the number of the projected populations, we took the nearest 24 cells and repeated this procedure until this criterion was met. Then the projected population and non-redundant hematopoietic cells were downsampled to an equal number of cells (maximum 500). For scATAC-seq, we binarized the matrix for both the projected populations and hematopoietic matrices. Next, we scaled the sparse matrices to 10,000 total counts for scRNA and 5,000 total promoter counts for scATAC-seq (promoter peaks defined as peaks within 500 bp of TSS from hg19 10x v3.0.0 gtf file). Next, we computed row-wise t-tests for each feature. We then calculated the FDR using p.adjust(method=”fdr”). We then computed the log2 mean and log2 fold changes for each feature. We chose these parameters based on Soneson et al., study comparing analytical methods for differential expression^45^. For scRNA-seq, differential expression was determined by FDR < 0.01 and absolute log2 fold changes greater than 0.5. For scRNA-seq, differential expression was determined by FDR < 0.05 and absolute log2 fold changes greater than 0.05.

To identify differential genes for bulk leukemia RNA-seq, we downsampled the gene counts to 10,000 counts randomly for 250 times. We then projected and used the above framework to resolve differential genes with log2 fold change > 3 and FDR < 0.01. We then removed genes that were differential in 33% or higher of the normal samples to attempt to capture biased genes. In addition, we further removed genes differential in 50% or higher of the leukemia samples. This filtering biases our identified malignant genes to those variable across the leukemic types vs conserved across all leukemic types. We then took the average malignancy for each remaining gene for each leukemic type and used the top 300 variable malignant genes across the leukemic types for heatmap and LSI. For computing differential LSI, we binarized each gene being malignant or not for the 300 variable malignant genes and computed the TF-IDF transform followed by SVD (LSI). We then visualized this in 2 dimensions using uwot’s implementation of UMAP (50 SVD dimensions, n_neighbors = 50, min_dist = 0.005).

#### Matching scATAC-scRNA-seq pairs using Seurat Canonical Correlation Analyses

We wanted to be able to integrate our epigenetic and transcriptomic data and built off of previous approaches for integration^10, 35^. We found the approach that worked best for our integrative analyses was using Seurat’s Canonical Correlation Analysis. We performed integration for each biological group separately because (1) it improved alignment accuracy and (2) required much less memory. First, for both the Gene Activity Scores matrix and scRNA matrix we created a Seurat Object “CreateSeuratObject”, then normalized with “NormalizeData”, and found the top 2000 variable genes/activities ranked by dispersion with “FindVariableGenes”. We then defined the union of the top 2000 variable genes from scRNA-seq and gene scores from scATAC-seq and found this increased the concordance downstream (defined by cluster to cluster mapping in hematopoiesis and single cell spearman correlations). These genes were then used for running Canonical Correlation analysis using “RunCCA” with the number of cc’s to compute as 25. We then calculated the explained variance using “CalcVarExpRatio” grouping by each of the individual experimental protocols scATAC (Gene Activity Scores) and scRNA. We then filtered cells where the variance explained by CCA is less than 2 fold compared to PCA. We then Aligned the subspaces with “AlignSubspace” and 25 dimensions to align with reduction.type = “cca” and grouping.var = “protocol”. We then identified for each scATAC cell the nearest scRNA cell based on minimizing the euclidean distance. We then created a UMAP using the aligned CCA coordinates as input into uwot’s UMAP implementation with n_neighbors = 50, min_dist = 0.5, metric = “euclidean” and then plotting with ggplot2 in R. To enable more robust correlation based downstream analyses, we used our initial KNN groupings (nGroups = 4998, KNN = 50) from Cicero^14^ to group scATAC accessibility, Gene Activity Scores, scRNA closest neighbor and chromVAR^15^ deviation scores.

#### Peak-To-Gene Linkage

Cicero^14^ allows us to infer Gene Activity Scores by linking distal correlated ATAC peaks to the promoter peak. While this measure is extremely useful, it does not actually mean it is correlated to the gene expression. To circumvent this limitation, we used our grouped scATAC and grouped linked scRNA-seq to identify peak-to-gene links. First we log-normalized the accessibility and gene expression with log2(Counts Per 10,000 + 1) and then we resized each of the gene GRanges to the start using resize(gr,1,”start”) and then resizing the start to a +- 250kb window using resize(gr, 2 * 250000 + 1, “center”). We then overlapped all ATAC-seq peaks using “findOverlaps” to identify all putative peak-to-gene links. We then split the aggregated ATAC and RNA matrices by whether majority of the cells were from MPAL or Hematopoietic single cells. We then correlated the peaks and genes for all putative peak-to-gene links. We used a previously described approach for computing a null correlation based on *trans* correlations (correlating peaks and genes not on the same chromosome)^2^. Briefly, for each chromosome 1000 peaks not on the same chromosome are identified and correlated to every gene on that chromosome. Each putative peak-to-gene correlation is converted into a z-score by using the mean and sd of the null *trans* correlations. These are then converted to p-values and adjusted for multiple hypothesis using the benjamini Hochberg correction “p.adjust” in R. We retained links whose correlation (Pearson) was above 0.35 and FDR < 0.1, same correlation cutoff as co-accessibility in Cicero^14^, in either MPAL or Hematopoietic aggregations. We then kept all peak-to-gene links that were greater than 2.5kb in distance. We identified peak-to-gene links that are only present in hematopoiesis, MPALs or both. To visualize the peak-to-gene links we plotted all of them as a heatmap with ComplexHeatmap. To determine the column order we first computed PCA for the first 25 PCs using irlba. We then computed Seurat^11^ Shared Nearest Neighbor clustering with a resolution of 1 and then computed the cluster means. We then computed the order of these clusters using hclust and the dissimilarity 1-R as the distance. Next we then iterated through each cluster and performed hclust with the dissimilarity calculations to get a final column order. The peak-to-gene links were grouped by k-means clustering with 10 input centers 100 iterations and 10 random starts for healthy, disease and the overlapping links. We did this bi-clustering because it enabled us to plot smaller rasterized chunks of the heatmap without overwhelming the memory and put the individual rasterized k-means clusters together post analysis.

#### Peak-To-Gene links enrichment with GTEx eQTLs

We adopted a previous approach for identifying the enrichment of our peak-to-gene links in GTEx eQTL data. Briefly, we downloaded GTEx eQTL data (version 7) from https://gtexportal.org/home/datasets and the *.signif_variant_gene_pairs.txt.gz files were used. We in addition downloaded gencode v19 (matched to these eQTLs) and identified all gene starts and identified all nearest gene starts to each peak and eQTL using “distanceToNearest”. We filtered all eQTLs that were further than 250kb from their predicted gene to be consistent with our linkage approach. To calculate a conservative overlap enrichment, we further pruned all eQTL links that were to its nearest gene. We then created a null set (n = 250) of peak-to-gene links by randomly selecting distal ATAC-seq peak-to-gene links (within 250 kb) that are distance matched to the links tested at 5kb resolution. We then calculated a z-score and enrichment for each peak-to-gene link set compared to the null set and calculated an FDR using p.adjust(method = “fdr”).

#### Peak-To-Gene links enrichment with K27ac HiChIP metaV4C

We wanted to determine the specify of our peak-to-gene links in published chromatin conformation data as previously described. We downloaded previously published Naive T cell and HCASMC K27ac HiChIP data. We then identified within each peak-to-gene links subset the peaks that were most biased to T/NK cells. To do this analysis, we calculated the z-score for each peak in the peak-to-gene links removed all links below 100kb and floored each peak coordinate (start or end) to its nearest 10kb window. We then ranked these links by the z-score for the peak, deduplicated the links at 10kb resolution and kept the top 500 remaining peak-to-gene links. Next, we used juicer dump (no normalization “NONE”) at 10kb resolution for each chromosome in the “.hic” file. Then we read each chromosomes into an individual “sparseMatrix” in R. We then scaled the sparse matrices such that the total cis interactions summed up to 10 million PETs. Then, for each peak-to-gene link, the upstream or downstream window (Column or Row) (whether the peak was upstream or downstream of the gene promoter) was identified. To scale each interactions distance for interpretability, we linearly interpolated the data to be on a −50-150% scale to visualize the focal interaction. The mean interaction signal was reported and repeated for both replicates. The mean and sd across both replicates were calculated and plotted with ggplot in R.

#### Identifying TF Malignant Target Genes and Survival Anlaysis

We wanted to create a framework for identifying TFs that potentially directly regulate malignant genes. To do this analysis, we first identified a set of transcription factors whose hypergeometric enrichment in differential peaks were high across the MPAL subpopulations (Comparing up-regulated peaks vs all peaks) and were identified as being transcriptionally correlated with their motif’s accessibility (see above). Next for a given TF and all identified peak-to-gene links, we further subsetted these links by those containing the TF motif. Then for each MPAL subpopulation, we determined for each peak-to-gene link if both the peak and gene are up-regulated. Then for each gene, we gave a binary score whether or not that MPAL subpopulation has at least one differential peak-to-gene link (whose peak and gene are differentially up-regulated) and report the proportion of subpopulations that were up-regulated. In addition, for each gene that has at least 1 differential peak-to-gene links we summed their squared correlation R^2^ and report that as the differential linkage score. We kept all genes that had least 1 MPAL subpopulation with corresponding differential peak-to-gene links.

For survival analysis, we downloaded the RPKM TCGA-LAML data^42^ (https://tcga-data.nci.nih.gov/docs/publications/laml_2012/laml.rnaseq.179_v1.0_gaf2.0_rpkm_matrix.txt.tcgaID.txt.gz). We downloaded the survival data from Bioconductor RTCGA.clinical (“patient.vital_status”) and matched using TCGA IDs the RPKM expression. Next, we took all genes that were identified as target genes for *RUNX1* (n = 732), and computed row-wise z-scores for each gene. Next, we took the column means of this matrix to get an average z-score across all *RUNX1* target genes. We then identified the top 33% and bottom 33% of donors based on this expression. We computed the p-value using the R package survival “survfit(Surv(times, patient.vital_status)∼Runx1_TG_Expression, LAML_Survival)”. We plotted the Kaplan-Meier curve using the R package survminer “ggsurvplot” in R.

## SUPPLEMENTARY FIGURE LEGENDS

**Supplementary Figure 1. Quality control of scRNA-seq data for hematopoiesis samples.**

**a**, (Top) Number of cells passing filter for each experimental replicate (number of informative genes > 400 and number of unique molecular identifiers (UMI) > 1000), (Middle) number of informative genes detected per single cell and (Bottom) number of unique molecular identified (UMI) transcripts.

**b**, Aggregated scRNA-seq one to one reproducibility plots for experimental replicates and across experiments.

**c**, scRNA-seq biological cluster labels assigned to each cluster overlay on UMAP of hematopoiesis.

**d**, scRNA-seq experimental sample labels overlay on UMAP of hematopoiesis.

**Supplementary Figure 2. Quality control of scATAC-seq data for hematopoiesis samples.**

**a,** scATAC-seq cell filtering plot. The x-axis is the number of unique accessible fragments and the y-axis is the enrichment of Tn5 insertions at transcription start sites, representing the robust signal to background for each single cell.

**b,** Aggregated scATAC-seq fragment size distributions across individual experiments demonstrating sub-, mono- and multi nucleosome spanning ATAC-seq fragments.

**c,** (Top) Number of cells passing filter for each experimental replicate (Unique fragments > 1000 and TSS enrichment > 8), (Middle) log10 unique fragments, (Middle) fraction of Tn5 insertions in the healthy hematopoietic union peak set, and (Bottom) enrichment at transcription start sites.

**d,** Aggregated scATAC-seq one to one reproducibility plots for experimental replicates and across experiments.

**e,** scATAC-seq biological cluster labels assigned to each cluster overlay on UMAP of hematopoiesis.

**f**, scATAC-seq experimental sample labels overlay on UMAP of hematopoiesis.

**Supplementary Figure 3. Quality control of scADT-seq data for hematopoiesis.**

**a,** Proportion of scRNA-seq cells passing filter that were matched with corresponding scADT data.

**b,** Aggregated scADT-seq one to one reproducibility plots for experimental replicates and across experiments.

**c,** scADT-seq UMAP of bmmc and pbmc samples across 14 antibodies. scADT overlay of experimental sample labels, *CD19*, *CD3*, *CD56*, *CD4*, *CD8A*, *CD14*, *CD16*, *CD45RA*, *CD45RO*, *TIGIT* and *PD-1*. Color represents experimental labels or scADT-seq values after CLR transformation.

**d,** Corresponding scRNA-seq biological cluster label overlay on the scADT-seq UMAP of BMMC and PBMCs.

**Supplementary Figure 4. Validation of key marker genes for both scRNA-seq and scATAC-seq for hematopoiesis.**

**a-h,** Multi-omic tracks; (Top) average track of all clusters displayed, (Middle) binarized 100 random scATAC-seq tracks for each locus at 100bp resolution and (right) scRNA-seq log2 distribution of normalized expression for each cluster, box-plot shows median and lower and upper quartiles.

**a,** Multi-omic track of *GATA1* (specific in these clusters for Erythroid) for erythroid development from HSC progenitor cells.

**b,** Multi-omic track of *GATA2* (specific in these clusters for Basophil) for erythroid development from HSC progenitor cells.

**c,** Multi-omic track of *ELANE* (specific in these clusters for GMP/Neutrophil) for neutrophil development from HSC progenitor cells.

**d,** Multi-omic track of *IRF8* (specific in these clusters for pDC) across pDC development from HSC progenitor cells.

**e,** Multi-omic track of *SDC1* (specific in these clusters for Plasma cells) across B cell development and plasma cells.

**f,** Multi-omic track of *CD1C* (specific in these clusters for cDC) across cDC development from HSC progenitor cells.

**g,** Multi-omic track of *SELL* (specific in these clusters for Naive T cells vs memory, and CD8 central memory vs CD8 effector memory) across NK and T cells.

**h,** Multi-omic track of *GZMB* (specific in these clusters for NK cells) across NK and T cells.

**Supplementary Figure 5. Diagnostic flow cytometry plots for MPALs 1-3.**

**a-c,** Diagnostic flow cytometry plots from three different MPAL cases gated on blasts area (highlighted in red) and lymphocytes (highlighted in black) from CD45 and side scatter area (SSC-A).

**a**, MPAL 1 shows classic bilineal phenotype with both T-lymphoblasts (cCD3-positive and CD7-positve) and myeloid blasts (MPO-positive and CD33-positive).

**b**, MPAL 2 demonstrates a more complex phenotype with both biphenotypic (single population expressing lymphoid marker CD7 and myeloid marker CD33) and bilineal T-Myeloid patterns (subpopulation expressing monocytic markers CD64, CD33, and CD14).

**c**, MPAL 3 demonstrates a classic biphenotypic case with coexpression of both T-lineage markers (cCD3-positive) and myeloid markers (MPO-positive).

**Supplementary Figure 6. Diagnostic flow cytometry plots for MPALs 4-5R.**

**a-c,** Diagnostic flow cytometry plots from three different MPAL cases gated on blasts area (highlighted in red) and lymphocytes (highlighted in black) from CD45 and side scatter area (SSC-A).

**a**, MPAL4 demonstrates a classic bilineal B/M phenotype expressing B-lineage markers (CD79a and CD19-positive) and myeloid markers (MPO-positive and CD33-positive).

**b**, MPAL5 demonstrates a more complicated phenotype with a subpopulation of blasts expressing T-lineage markers (cCD3-positive and CD7-positive) and a subpopulation expressing myeloid marker MPO.

**c**, MPAL5R post-treatment relapse of MPAL5. Flow cytometry reveals expansion of the T-lymphoblastic subpopulation (cCD3-positive, TdT-positive population) following chemotherapy.

**d**, High-confidence mutations detected in 5 MPAL cases by whole exome sequencing. Missense mutations are shown in blue, frameshift deletions are shown in yellow, stopgain mutations are shown in purple, frameshift insertions are shown in orange, and nonframeshift deletions are shown in dark gray.

**Supplementary Figure 7. Quality control of scRNA-seq and scATAC-seq data for MPAL samples.**

**a,** (Top) Number of cells passing filter for each experimental replicate (number of informative genes > 400 and number of unique molecular identifiers (UMI) > 1000), (Middle) number of informative genes detected per single cell and (Bottom) number of unique molecular identified (UMI) transcripts.

**b,** Aggregated scRNA-seq one to one reproducibility plots for experimental replicates and across experiments.

**c,** scATAC-seq cell filtering plot. The x-axis is the number of unique accessible fragments and the y-axis is the enrichment of Tn5 insertions at transcription start sites, representing the robust signal to background for each single cell.

**d,** Aggregated scATAC-seq fragment size distributions across individual experiments demonstrating sub-, mono- and multi nucleosome spanning ATAC-seq fragments.

**e,** (Top) Number of cells passing filter for each experimental replicate (Unique fragments > 1000 and TSS enrichment > 8), (Middle) log10 unique fragments, (Middle) fraction of Tn5 insertions in the MPAL union peak set, and (Bottom) enrichment at transcription start sites

**f,** Aggregated scATAC-seq one to one reproducibility plots for experimental replicates and across experiments.

**Supplementary Figure 8. Evaluation of LSI projection workflow for previously published bulk and single-cell hematopoietic data sets across different platforms.**

**a,** Overview of LSI projection workflow. Briefly, using information from TF-IDF transform, singular value decomposition and UMAP of hematopoiesis enables projection of new data into the same subspace.

**b**, LSI projection of downsampled previously published bulk sorted hematopoietic data sets^18, 20^. (Left) RNA-seq downsampled bulk projections for 49 samples (n=250 downsampled cells). (Right) ATAC-seq downsampled bulk projections for 90 samples (n=250 downsampled cells).

**c**, LSI projection of downsampled previously published single-cell hematopoietic data sets labeled by previous classifications^20–22^. (Left) scRNA-seq projections of previous study healthy bone marrow cells (different platform and different aligned genome) colored by previous classifications. (Right) scATAC-seq projections for healthy bone marrow and peripheral blood samples (2 different platforms across 3 studies), colored by ground truth isolated populations.

**Supplementary Figure 9. LSI projection of previously published healthy and AML scRNA-seq identifies malignant programs across AML subpopulations.**

**a**, (Left) Schematic of LSI projection. (Right) Initial projection of all AML malignant single-cells colored by previous classifications^19^.

**b**, Re-classification of scRNA-seq AML single-cells based on closest normal cells in healthy hematopoiesis (See Methods). Broader re-classification increases the number of cells per category for improved power in differential analyses. LSI projection for each individual AML samples onto scRNA-seq healthy hematopoiesis colored by re-classifications (denoted is the sample id and number of cells).

**c**, K-means differential scRNA-seq heatmap (k = 10), colored by log2 fold change, comparing each AML sample subpopulations (classifications) vs their closest normal bone marrow cells from the same study^19^.

**Supplementary Figure 10. scADT-seq overlay of MPALs projected onto the hematopoietic hierarchy**

**a**, (Left) Projected MPALs colored by hematopoietic compartments. (Right) scADT-seq overlay of *CD7*, *CD33*, *CD14*, *CD4 and CD19* on MPAL single cells LSI projected onto hematopoiesis.

**Supplementary Figure 11. Visualization of differential genes and accessible peak regions.**

**a**, Top conserved differential genes across the MPAL hematopoietic compartments.

**b**, Top conserved differential transcription factors across the MPAL hematopoietic compartments.

**c**, KEGG pathway enrichment in differential RNA k-means 2, 3, 4, and 10 (Figure 2c).

**d-e,** Multi-omic differential tracks (Left) scATAC tracks showing MPAL disease subpopulations (red) closest normal cells (grey). (Right) Distribution of log2 normalized expression for MPAL disease subpopulations (red) and closest normal cells (grey); black line represents the mean and asterisk denote significance (LFC > 0.5 and FDR < 0.01).

**d,** Multi-omic differential track of *CDK11A*, up-regulated in MPALs 1, 2, 5 and 5R.

**e,** Multi-omic differential track of *CDKN2A*, up-regulated in MPALs 1, 2, 3, 4, and 5.

**Supplementary Figure 12. Seurat canonical correlation analysis alignment of scRNA and scATAC-seq hematopoietic and MPAL samples.**

**a**, Schematic of LSI projection of downsampled bulk leukemia RNA-seq onto healthy hematopoiesis.

**b**, Representative downsampled LSI projections (n=250) for B-ALLs, non-ETP T-ALLs, ETP T-ALLs, AMLs, T/M MPALs and B/M MPALs from previous studies^16^.

**c**, LSI UMAP of differentially up-regulated gene expression profiles across bulk leukemias^16^ and MPAL samples assayed in this study, colored by cytogenetics.

**d**, Binary heatmap of variable malignant genes across leukemia classifications. Each cell in the heatmap is colored whether the gene was identified as malignant for the leukemic sample.

**Supplementary Figure 13. Seurat canonical correlation analysis alignment of scRNA and scATAC-seq hematopoietic and MPAL samples.**

**a**, UMAP of CCA alignment of scATAC-seq using Cicero gene activity scores and scRNA-seq for (Left) bone marrow, (Middle) CD34+ enriched bone marrow, (Right) peripheral blood.

**b**, UMAP of CCA alignment of scATAC-seq using Cicero gene activity scores and scRNA-seq for MPAL samples.

**Supplementary Figure 14. Evaluation of scRNA and scATAC-seq alignment and peak-to-gene linkage across hematopoiesis and MPAL samples.**

**a**, Spearman rank correlation between scATAC-seq Cicero gene activity scores to scRNA-seq for each mapped cell within across all biological experiments.

**b**, Pearson correlation of CCA scRNA and scATAC-seq nearest-neighbors. The cutoff (R > 0.45) for high quality nearest neighbor mappings is shown.

**c**, (Left) UMAP of scATAC-seq hematopoiesis colored by scATAC-seq clusters. (Right) UMAP of scATAC-seq hematopoiesis colored by mapped scRNA-seq clusters.

**d**, Confusion matrix of initial clusters for mapped scRNA-seq to scATAC-seq clusters for hematopoiesis (Figure 1b-c).

**e**, (Left) Distribution of peak-to-gene distances. (Left-Middle) Distribution of number of peaks mapped per gene (median = 6). (Right-Middle) Distribution of number of genes mapped per peak (median = 1). (Right) Distribution of number of genes skipped for peak-to-gene links (median = 2).

**f**, MetaV4C plots of K27ac HiChIP in Naive T and HCASMC cells for top 500 biased T/NK (broad classification) peak-to-gene links that are identified only in healthy hematopoiesis. Shading indicates standard deviation between replicate experiments (n = 2).

**g**, Peak-to-genes enrichment in GTEx eQTLs over a permuted background distance-matched set (n=250) for the union set of peak-to-gene links.

**Supplementary Figure 15. Peak-to-gene links nominate putative regulatory regions that nominate key leukemic genes.**

**a-d,** Multi-omic differential track; (Middle) Aggregated scATAC tracks showing MPAL disease subpopulations (red) and closest normal cells (grey). (Right) Distribution of log2 normalized expression of gene of interest for MPAL disease subpopulations (red) and closest normal cells (grey); black line represents the mean and asterisk denote significance (LFC > 0.5 and FDR < 0.01). (Bottom) Peak-to-gene links for gene of interest.

**a**, Multi-omic differential track for *IL1RAP*.

**b**, Multi-omic differential track for *CD96*.

**c**, Multi-omic differential track for *FLT3*.

**d**, Multi-omic differential track for *MCL1*.

**Supplementary Figure 16. Analysis workflows for processing of scRNA-seq and scATAC-seq data.**

**a**, scRNA-seq analysis workflow. Briefly cells are aligned using 10x cell ranger, quality filtered, and clustered using a feature optimization approach (see methods).

**b**, scATAC-seq analysis workflow. Briefly cells are aligned using 10x cell ranger atac, quality filtered, clustered in large windows genome-wide, peak-calling on clusters, creation of a counts matrix and clustered using a feature optimization approach (see methods).

**Supplementary Table 1. MPAL Patient Characteristics.**

MPAL patient WHO Diagnosis, Age, Sex, Blast %, White Blood Cell Count, Cytogenetics, Prior Treatment.

**Supplementary Table 2. Antibodies used in flow cytometry of MPALs.**

**Supplementary Table 3. CITE-Seq Antibody List and Barcodes.**

Antibody information for Hematopoietic and MPAL samples. Barcodes used for sequencing ADT libraries.

**Supplementary Table 4. Differential analyses for MPAL and AMLs.**

MPAL differential RNA-seq k-means, MPAL differential ATAC-seq k-means, AML differential RNA-seq k-means and MPAL vs AML comparison.

**Supplementary Table 5. Motif enrichment and linkage to target genes.**

MPAL differential ATAC-seq k-means enrichment for CIS-BP motifs shown in figure 3A, all motifs, significant peak-to-gene links, and RUNX1 target genes.

## ACKNOWLEDGEMENTS

We thank M. Ryan Corces, Ansuman T. Satpathy and other members of the Chang and Greenleaf laboratories for helpful discussions. We thank the following people at 10x Genomics: Darisha Jhutty, Julia Lau, Josephine Lee, Luz Montesclaros, Katie Pfeiffer, Jessica Terry, Jean Wang, Yifeng Yin, and Solongo Ziraldo for help with sample preparation and library generation of scATAC and feature barcoding libraries. We acknowledge the Stanford Hematology Division Tissue Bank for providing samples for this study. Supported by the Swedish Research Council (grant 2015–06403, A.M.). Supported by P50-HG007735 (H.Y.C., W.J.G.), R35-CA209919 (H.Y.C.), Ludwig Cancer Research(R.M., H.Y.C.). H.Y.C. is an Investigator of the Howard Hughes Medical Institute. W.J.G is a Chan Zuckerberg Investigator.

## CODE AVAILABILITY

Code used in this study will be posted on GitHub for main analyses.

## DATA AVAILABILITY

Sequencing data will be deposited in the Gene Expression Omnibus (GEO). There are no restrictions on data availability or use.

## Author Contributions

L.M.M. and S.K. conceived the project and designed the experiments. L.M.M., M.L., E.G., and R.M. curated patient samples. S.K. led data production and performed the experiments together with A.K., A.M., and L.M.M.. G.X.Y.Z. provided healthy bone marrow and peripheral blood CITE-seq data. S.K. analyzed the scADT-seq data with contribution from B.P.. J.M.G conceived the analytical workflows and performed the data analysis for scATAC-seq and scRNA-seq supervised by H.Y.C. and W.J.G.. J.M.G., S.K., L.M.M., and W.J.G wrote the manuscript with input from all authors.

