## Supplementary Figures for "A single cell framework for multi-omic analysis of disease identifies malignant regulatory signatures in mixed phenotype acute leukemia"

**a**, Multi-omic differential track for *IL1RAP*.

**b**, Multi-omic differential track for *CD96*.

**c**, Multi-omic differential track for *FLT3*.

**d**, Multi-omic differential track for *MCL1*.

**Supplementary Figure 16. Analysis workflows for processing of scRNA-seq and scATAC-seq data.**

**a**, scRNA-seq analysis workflow. Briefly cells are aligned using 10x cell ranger, quality filtered, and clustered using a feature optimization approach (see methods).

**b**, scATAC-seq analysis workflow. Briefly cells are aligned using 10x cell ranger atac, quality filtered, clustered in large windows genome-wide, peak-calling on clusters, creation of a counts matrix and clustered using a feature optimization approach (see methods).

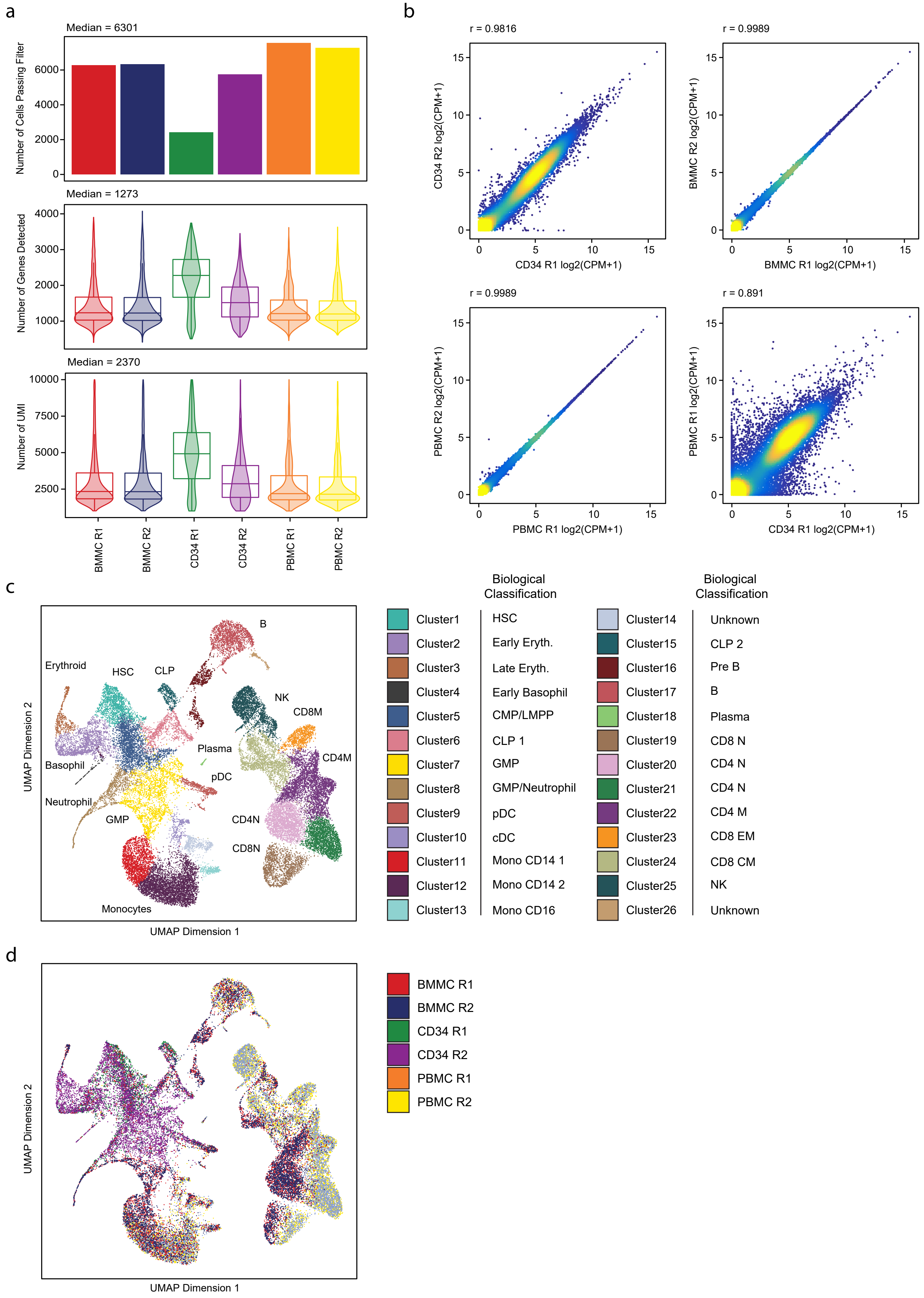


Supplemental Figure 1

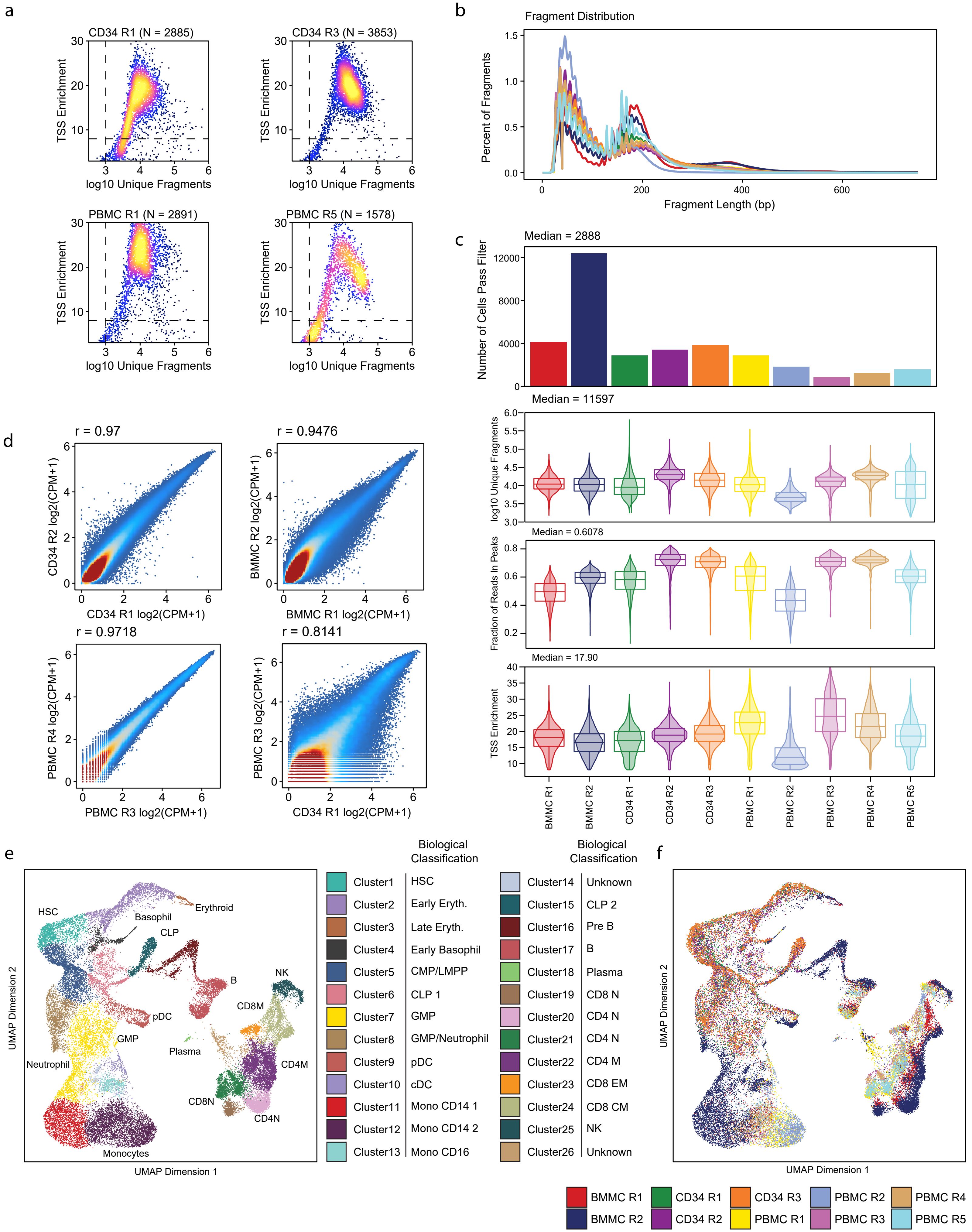


Supplemental Figure 2

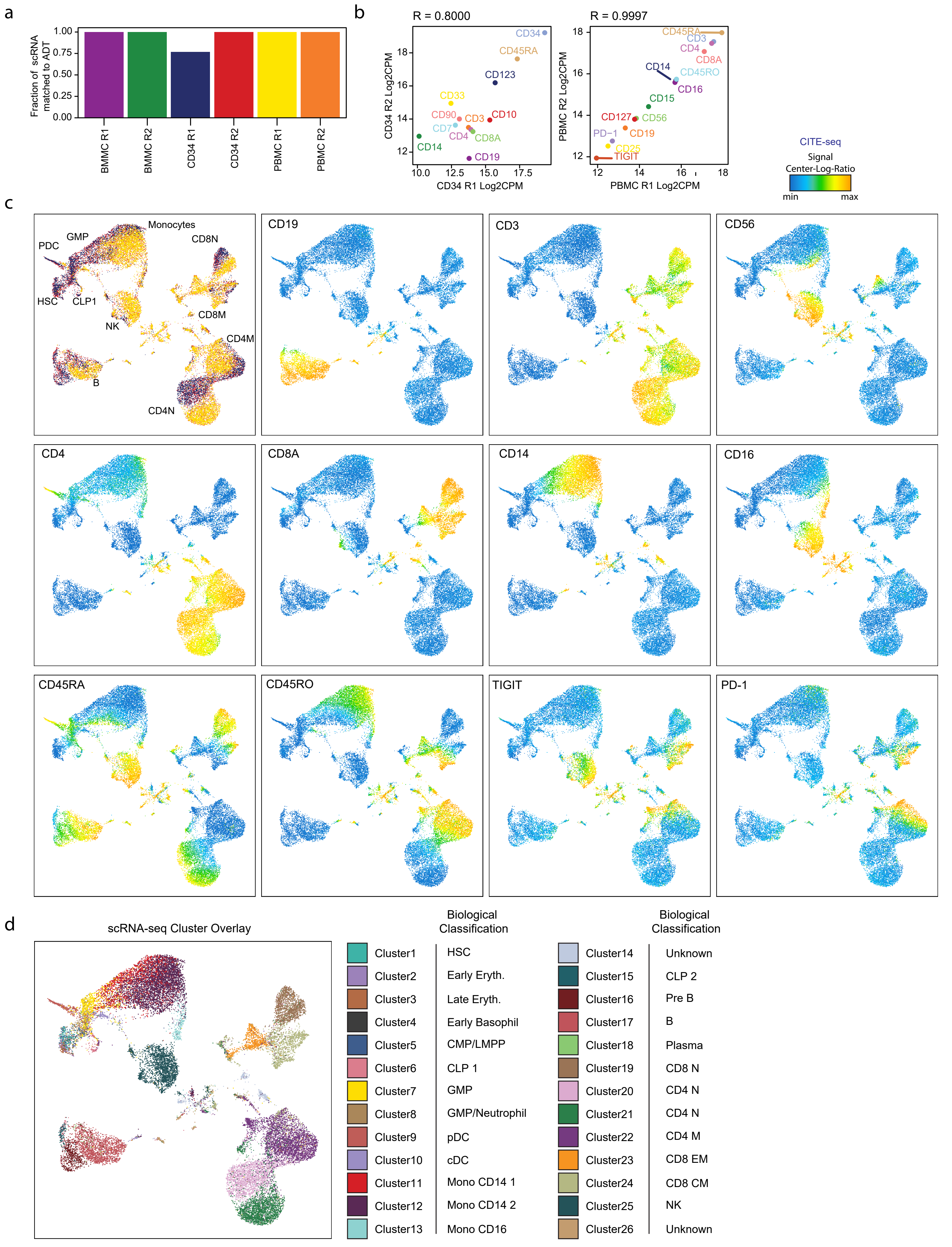


Supplemental Figure 3

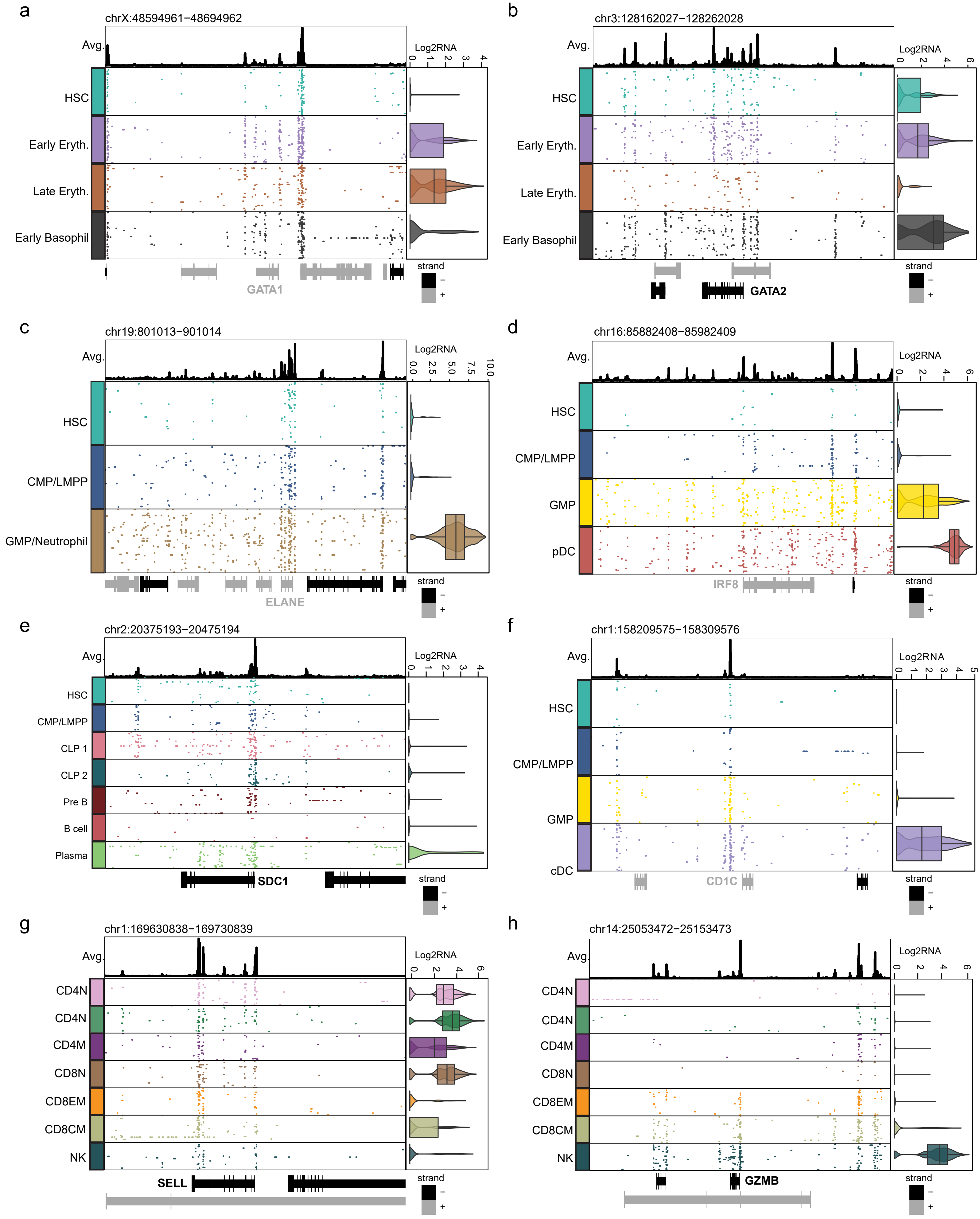


Supplemental Figure 4

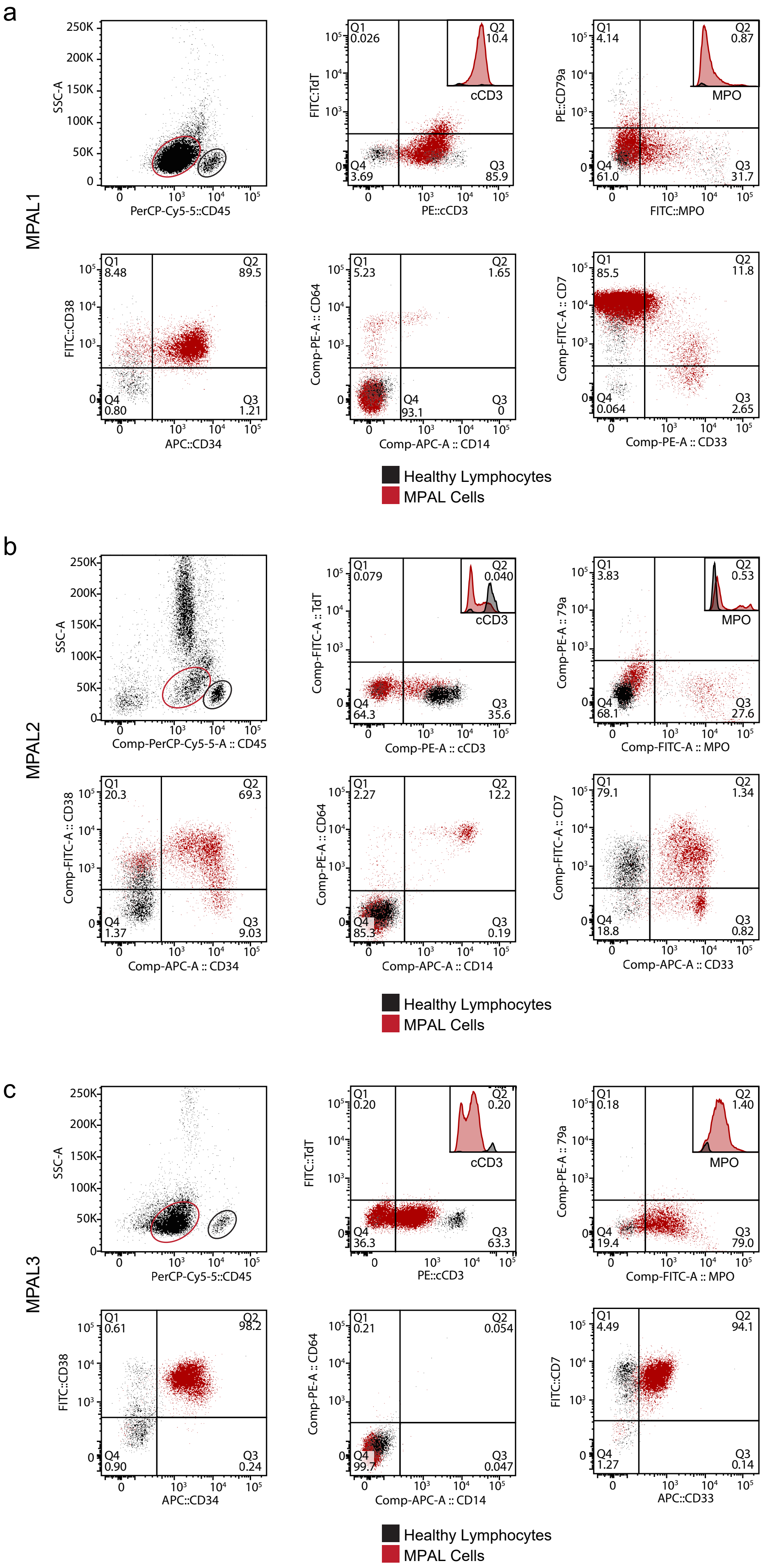


Supplemental Figure 5

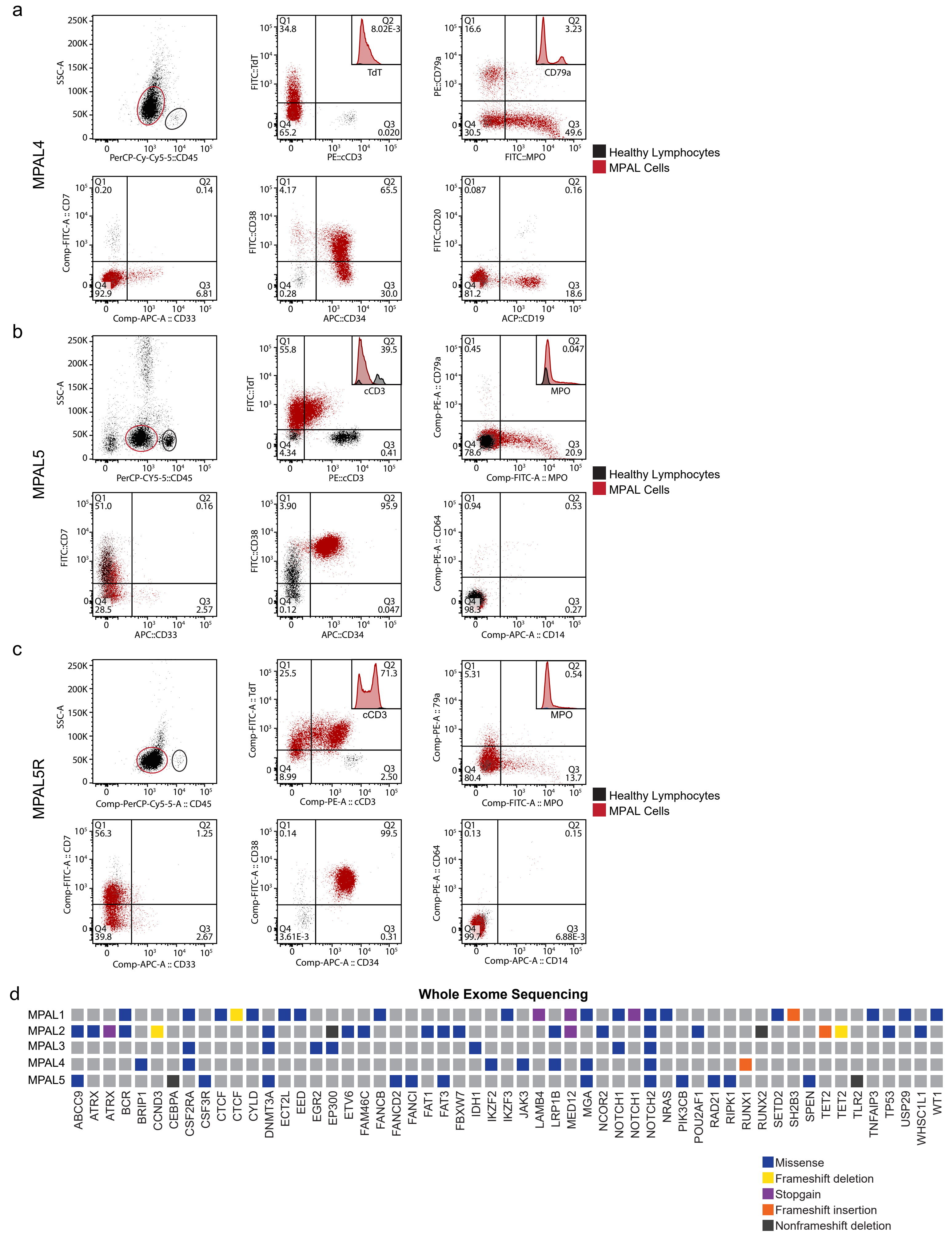


Supplemental Figure 6

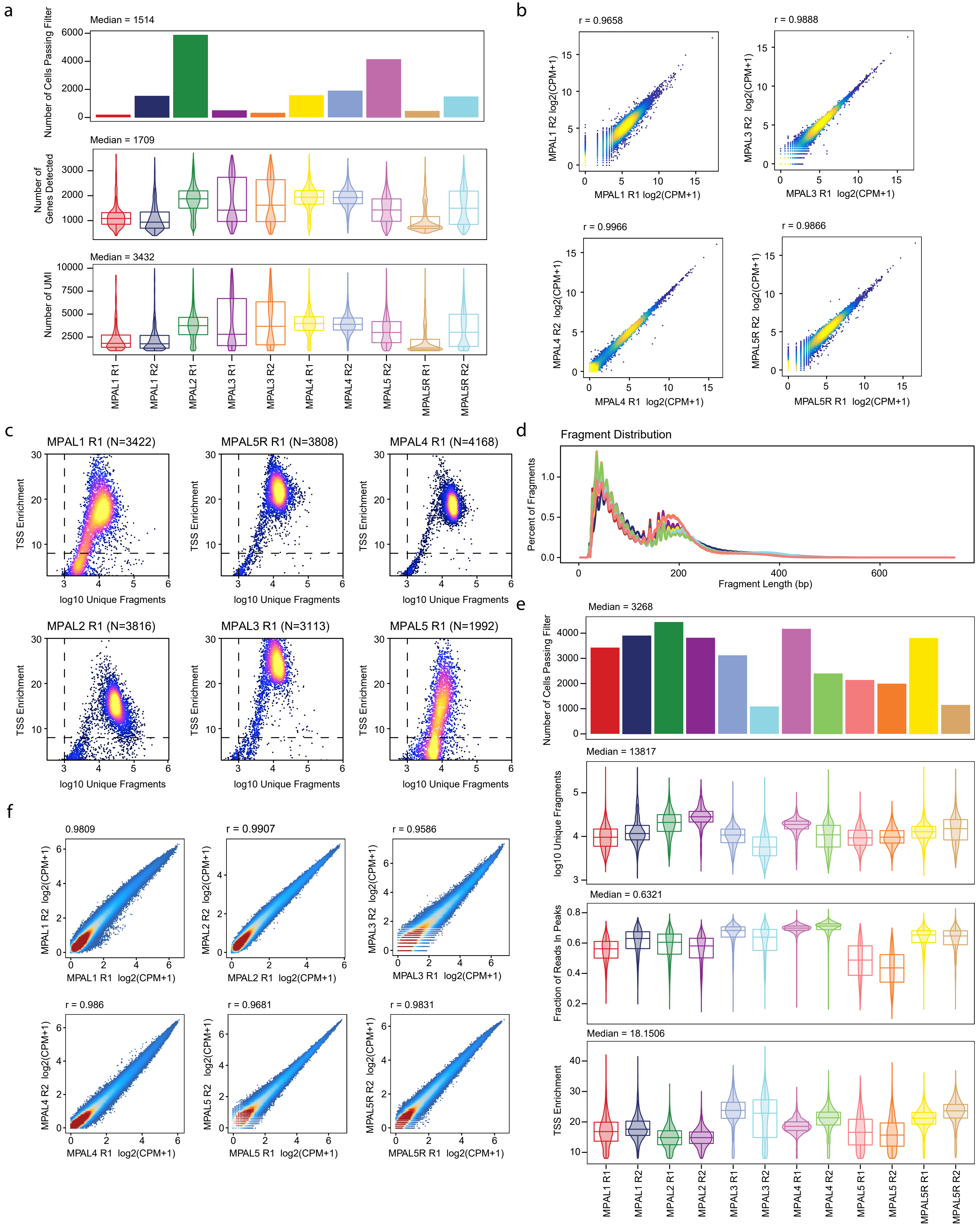


Supplemental Figure 7

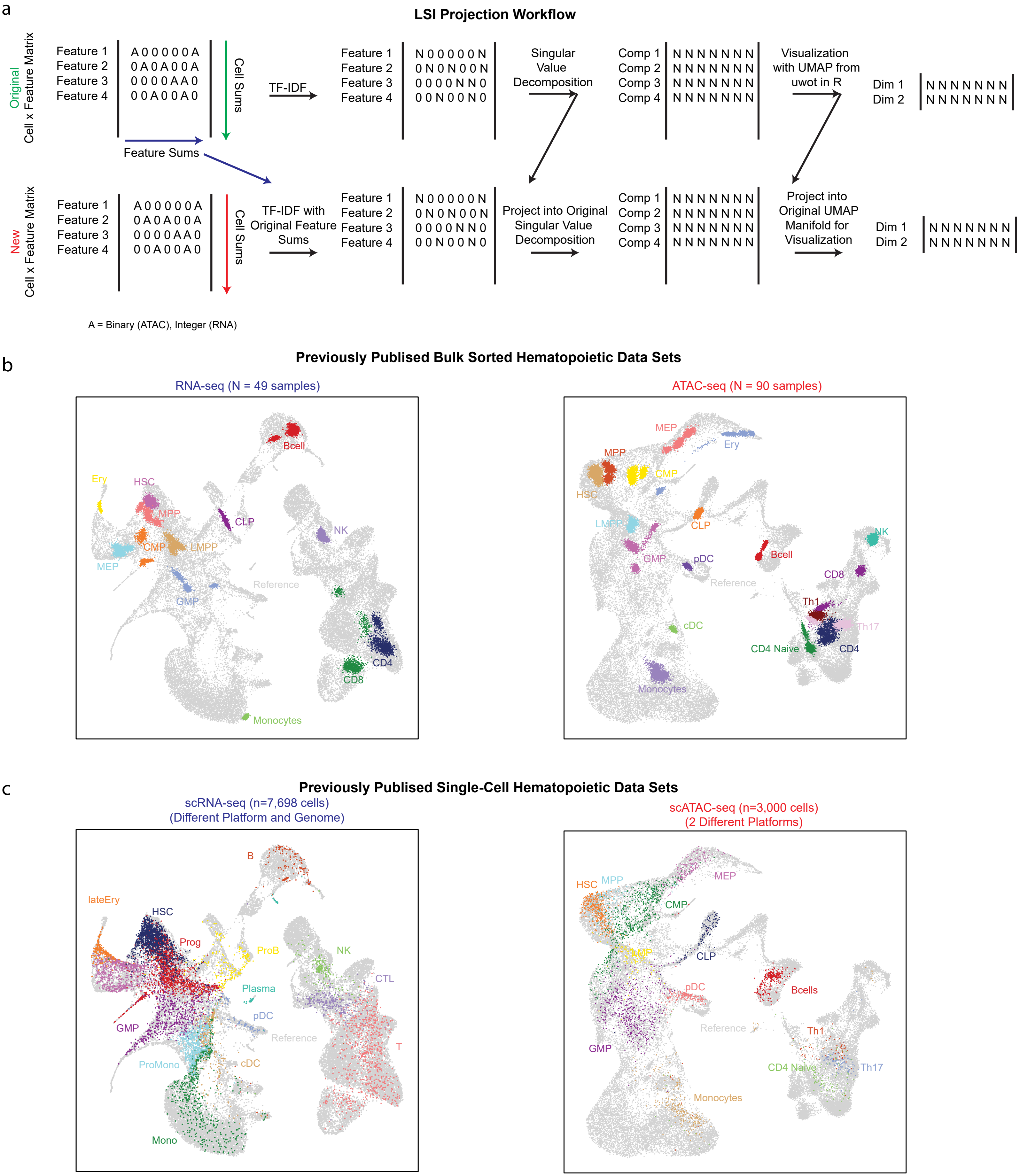


Supplemental Figure 8

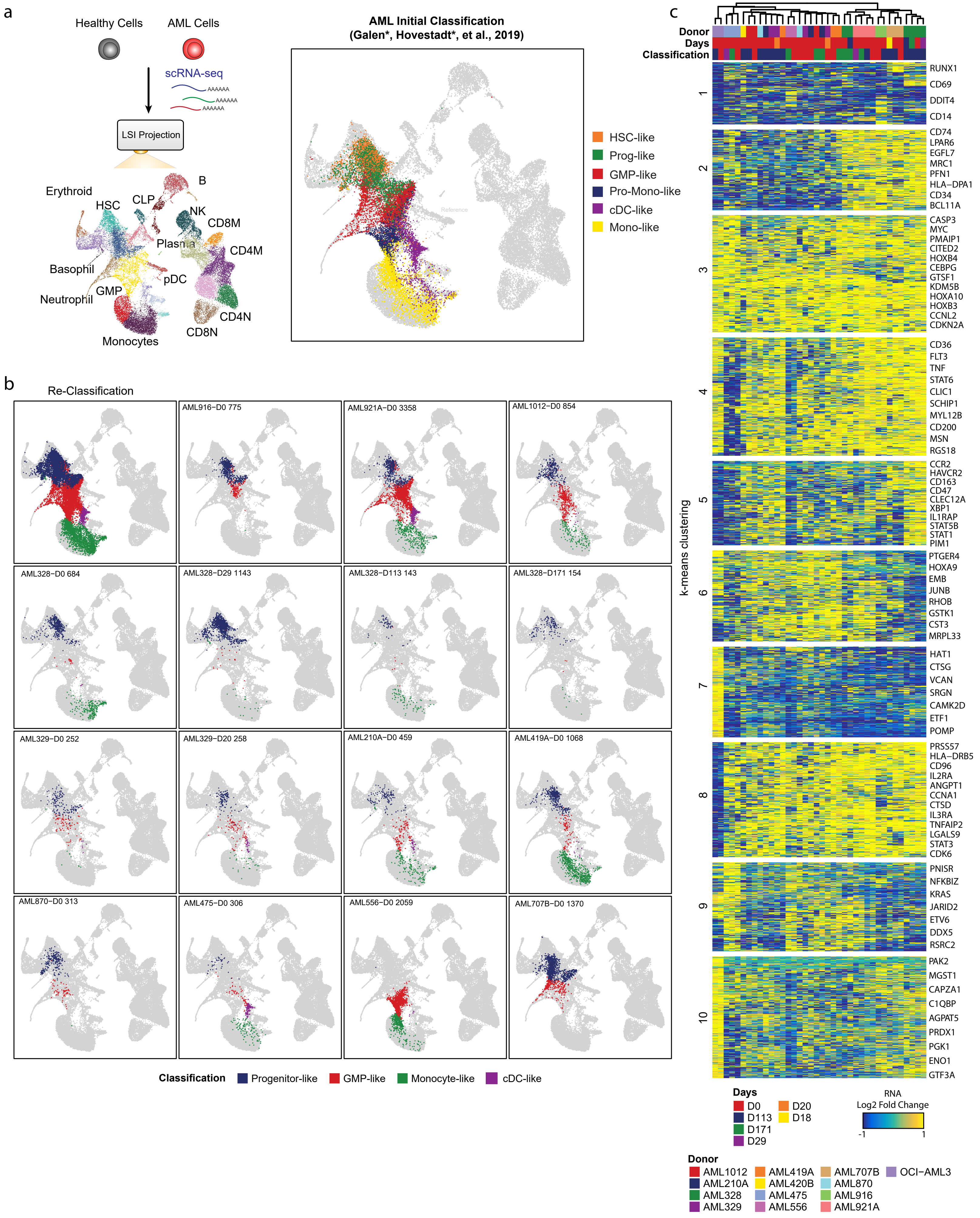


Supplemental Figure 9

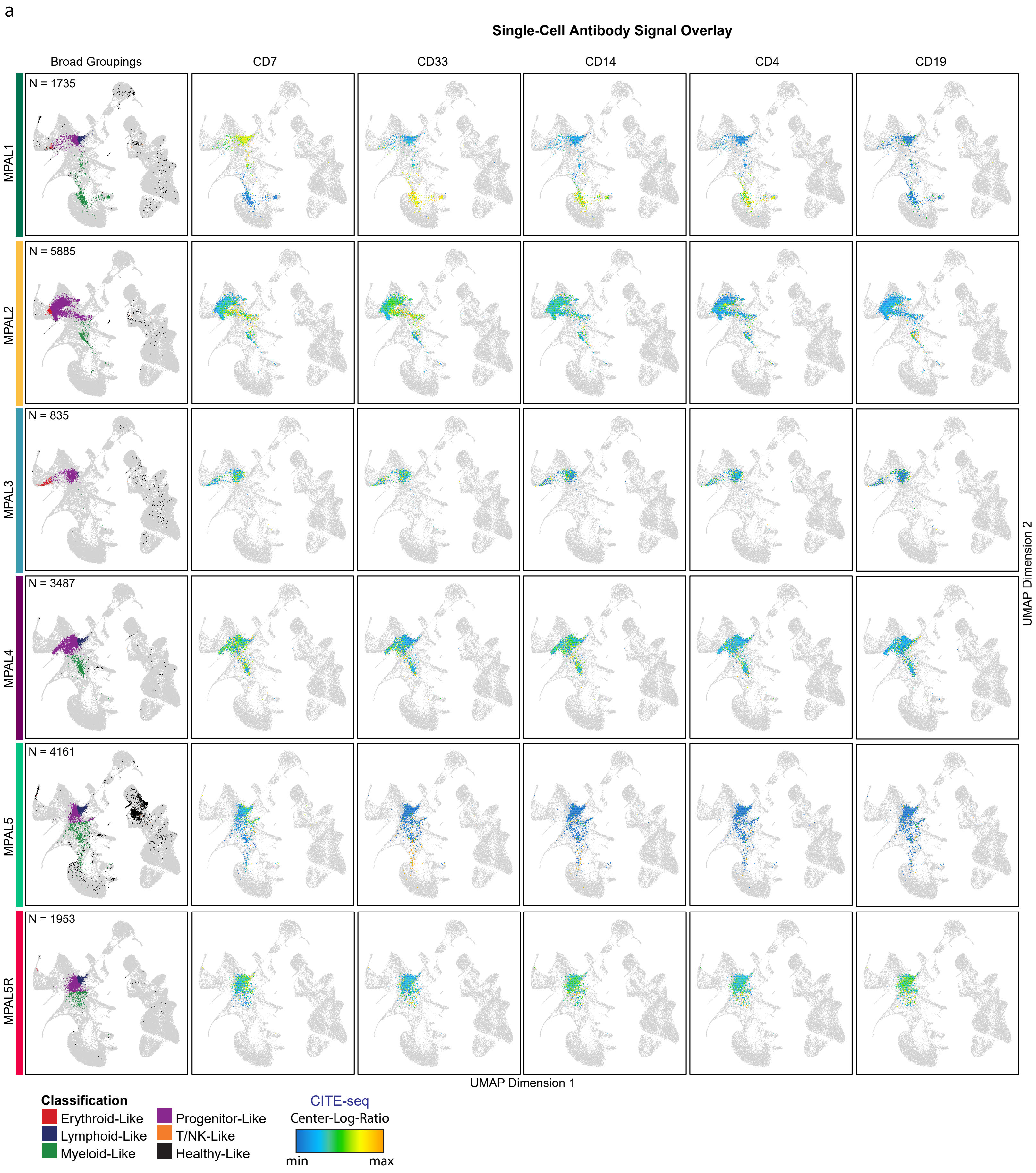


Supplementary Figure 10

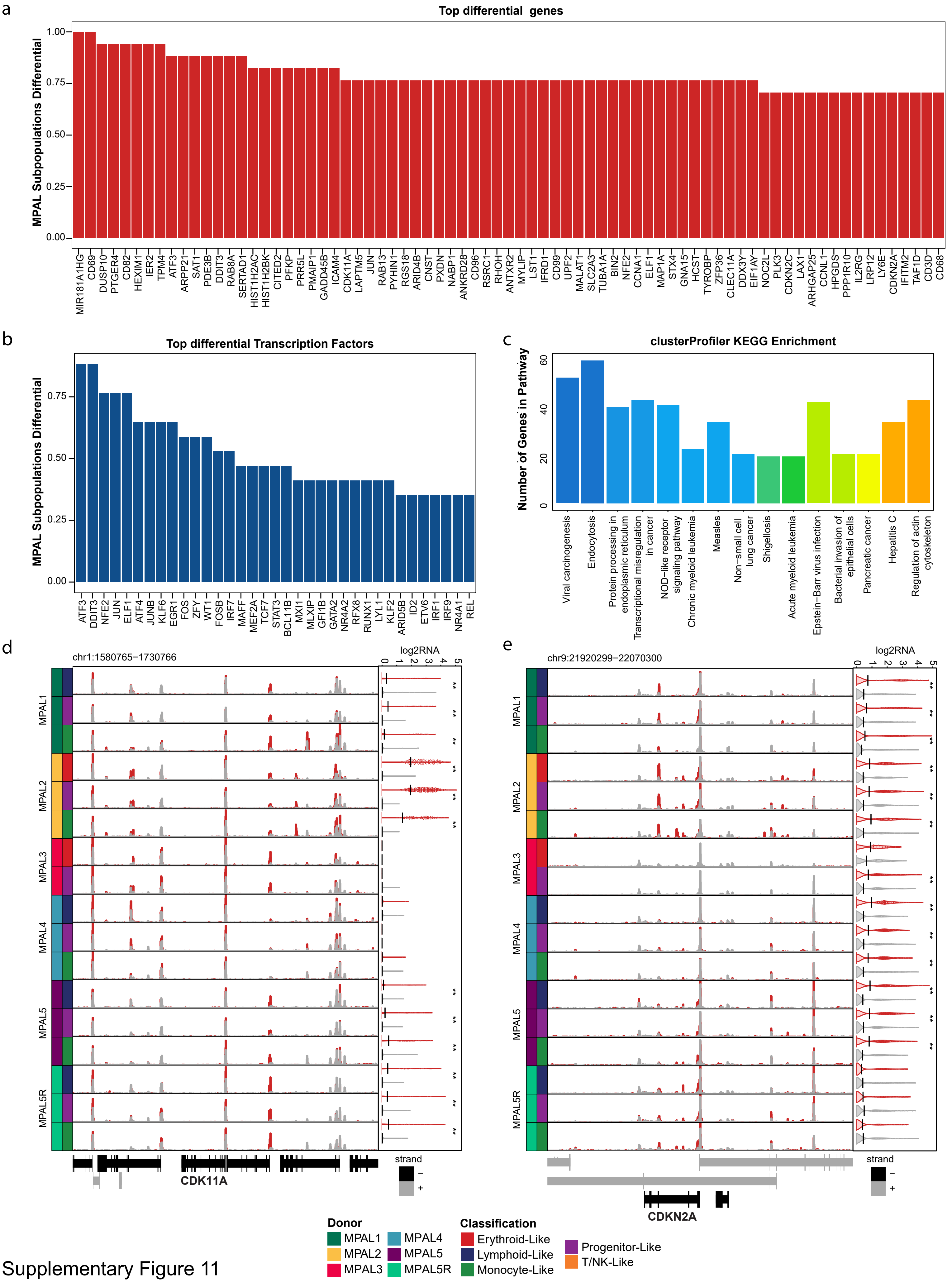


Supplementary Figure 11

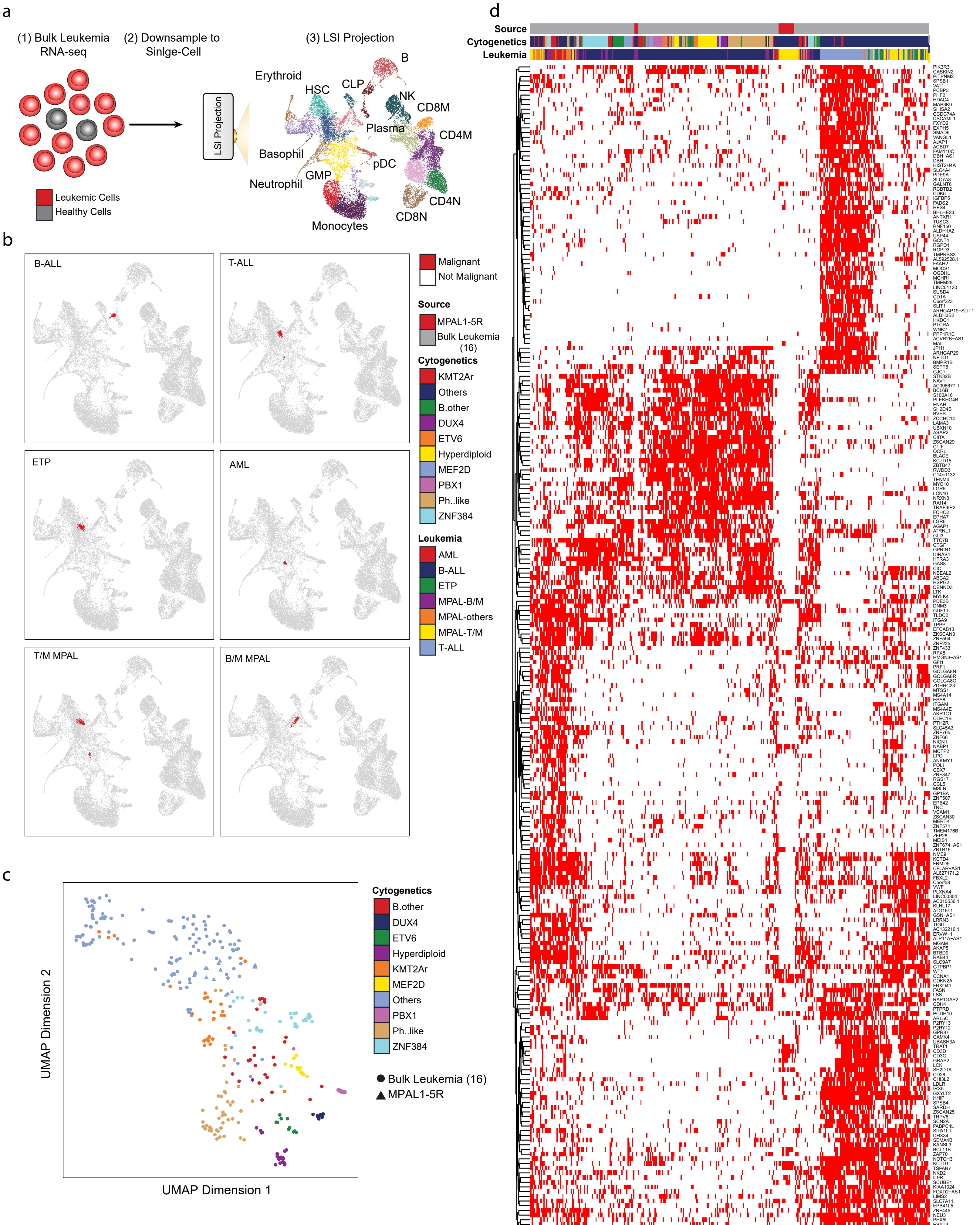


Supplementary Figure 12

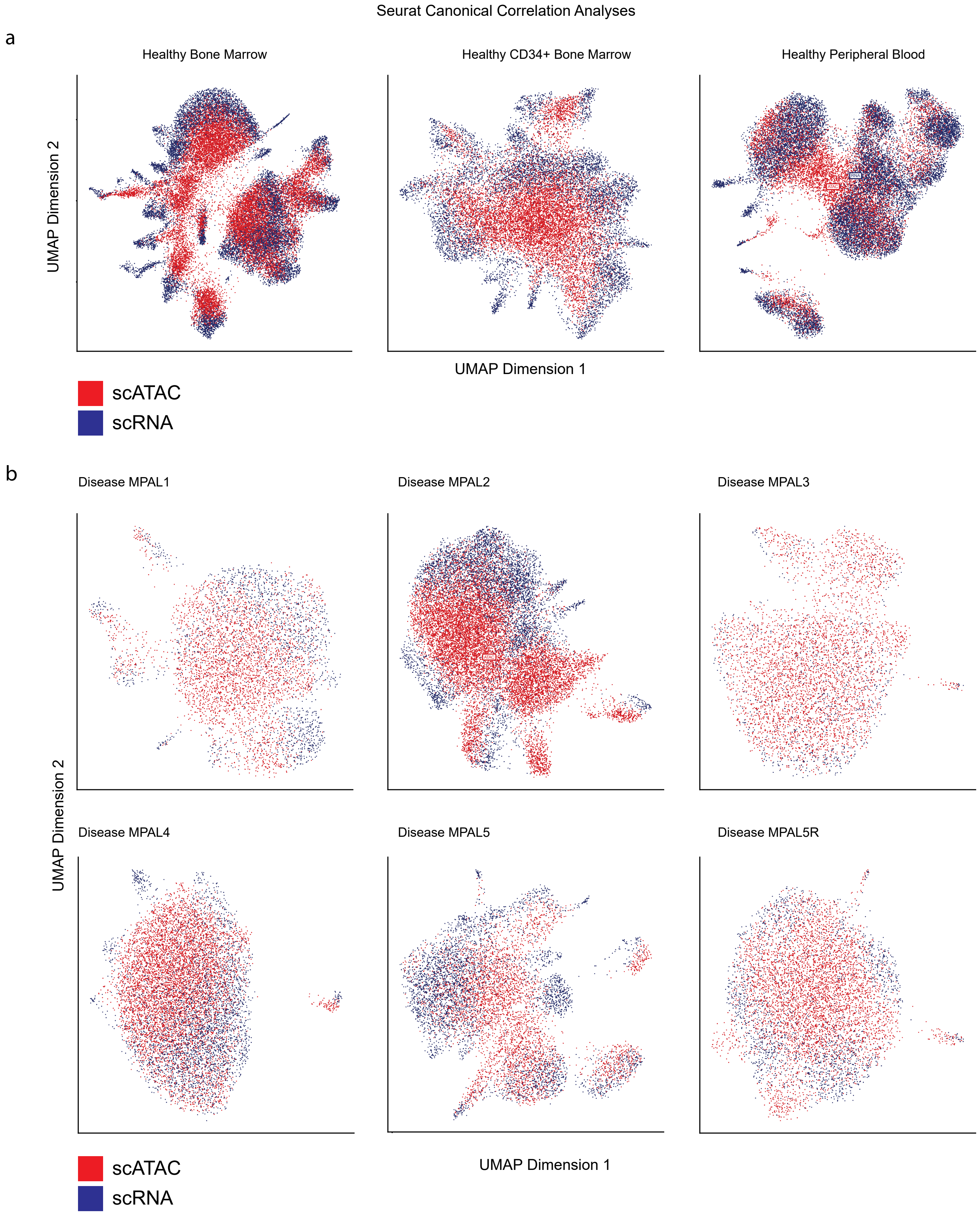


Supplemental Figure 13

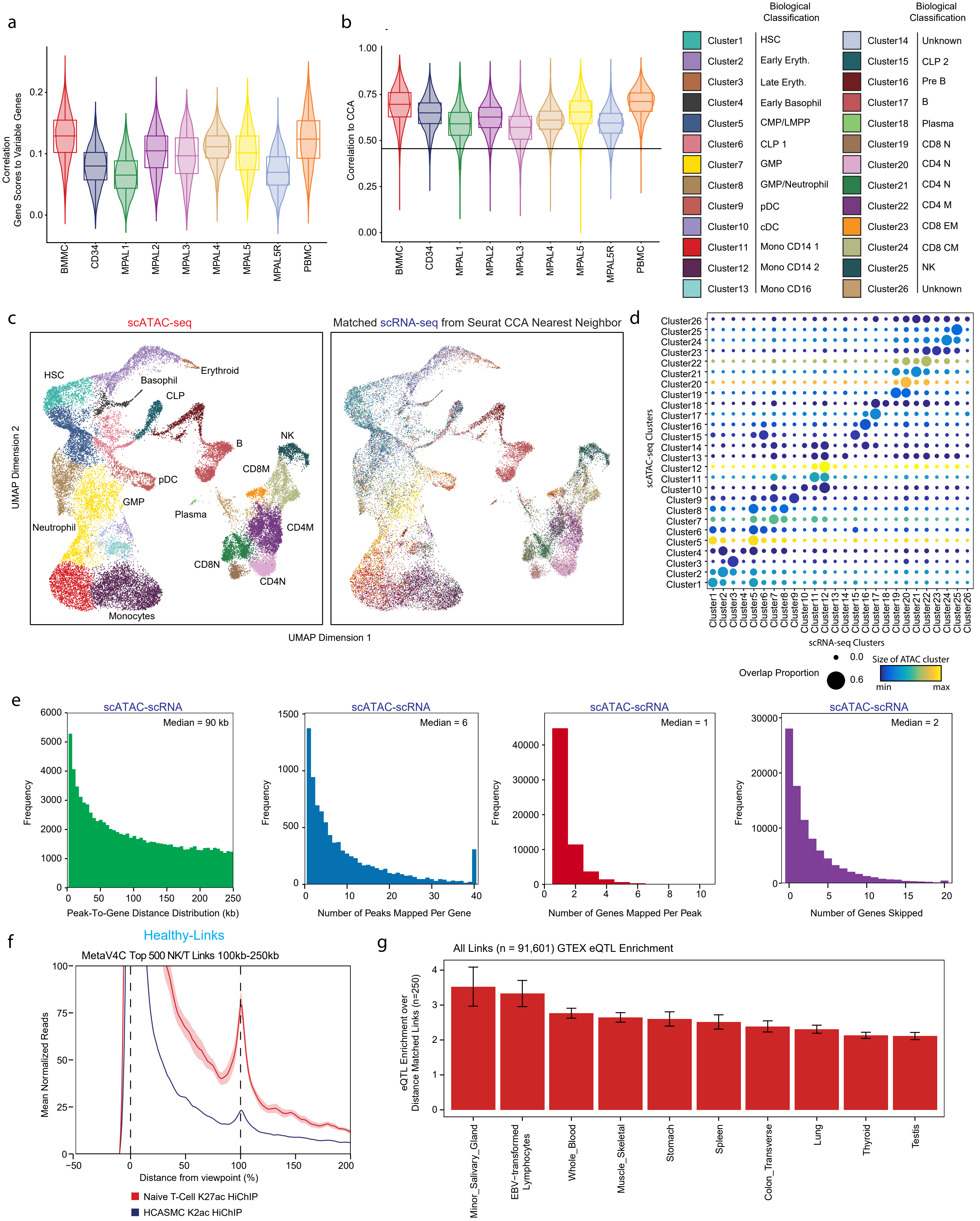


Supplementary Figure 14

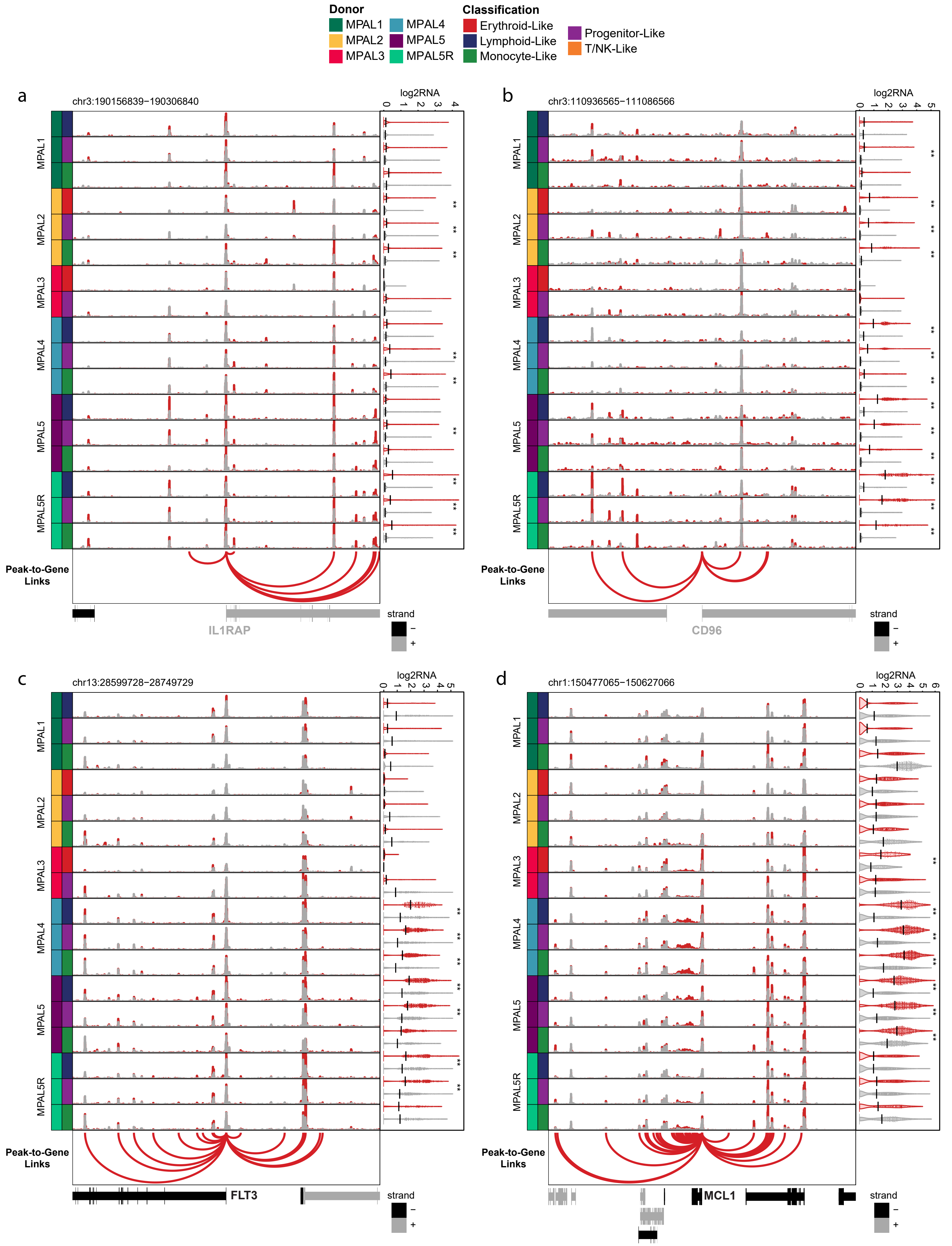


Supplementary Figure 15

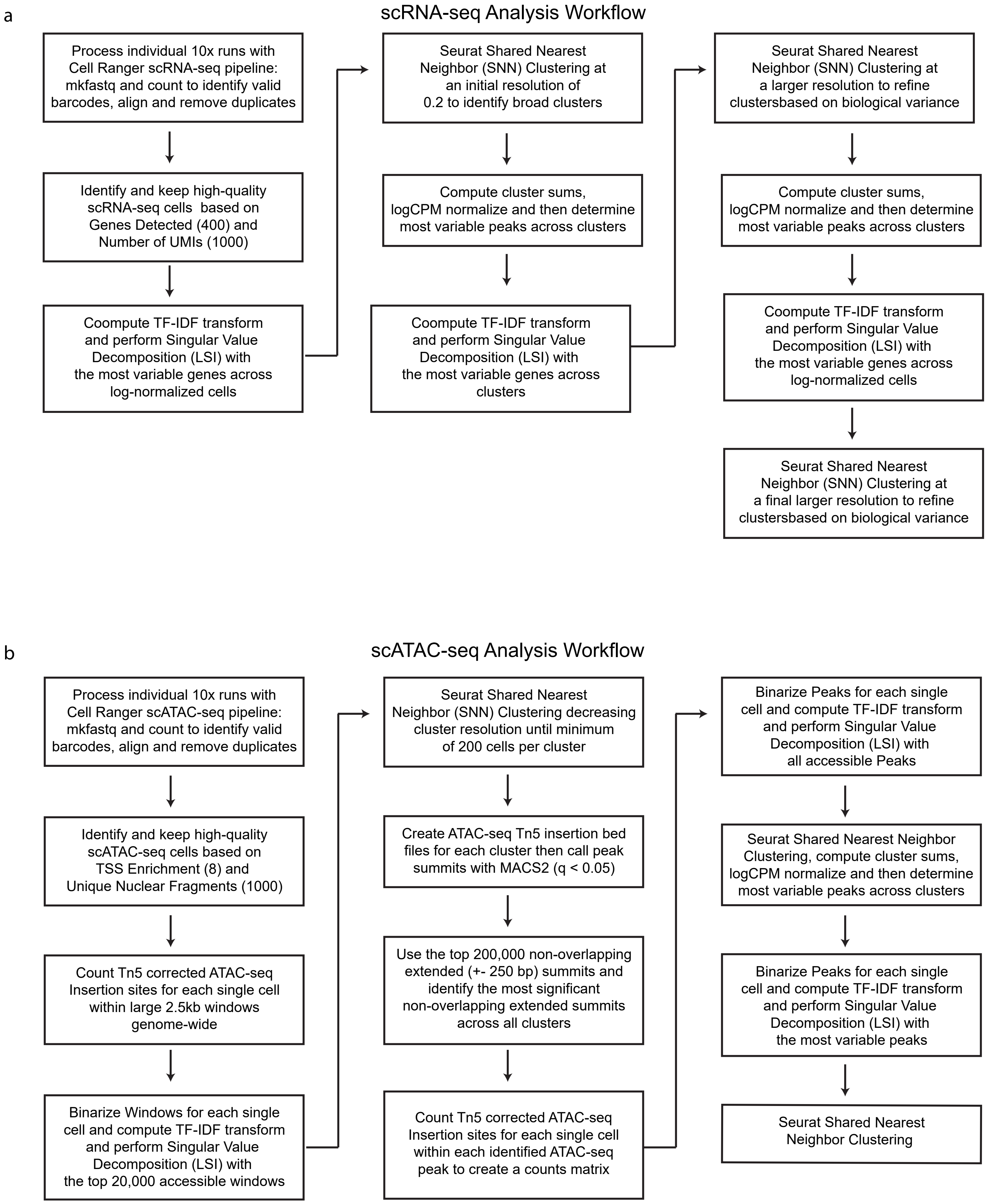


Supplemental Figure 16
